# The Training Village: an open platform for continuous testing of rodents in cognitive tasks

**DOI:** 10.64898/2026.01.12.698970

**Authors:** Balma Serrano-Porcar, Rafael Marin-Campos, Javier Rodríguez, Caterina Barezzi, Harshkumar Vasoya, Donna Kean, Duncan Pottinger, Alex Taylor, Hernando Martínez Vergara, Jaime de la Rocha

## Abstract

The development of new methods for dissecting neural circuits in rodents promises to transform the study of higher-order cognitive functions. However, the adaptation of complex cognitive tasks from humans to rodents remains very limited as these tasks require long periods of animal training with significant researcher expertise and effort. Multi-box setups can mitigate the problem but remain labor-intensive and costly to maintain. Newly developed home cage training systems, while minimizing costs, are limited to specific conditions and tasks. Here, we present the Training Village (TV), an open-source, affordable, and fully automated system for continuous rodent training. In the TV, a group of rodents lives in enriched arenas while individually accessing an operant box at any time. The system provides personalized training, continuous monitoring of task performance and home cage activity, and real-time remote supervision. Its graphical interface and modular design make it user-friendly and easy to integrate with other behavioral systems running a wide range of cognitive paradigms. We validated the TV across multiple tasks and cohorts of mice and rats, demonstrating efficient operant box usage, stable long-term task engagement, and its potential to link home cage behavior with task performance. Overall, the TV is an easy-to-adopt platform that can greatly accelerate brain research by fully automating rodent training in complex cognitive tasks.

## Introduction

The study of neural mechanisms underlying cognitive functions is expanding from humans to rodents, as cutting-edge tools and genetic models for dissecting neural circuits are largely restricted to these species^1–8^. However, adapting complex cognitive tasks to rodents remains challenging. It typically requires prolonged training periods with significant researcher involvement, including conducting repeated behavioral sessions with many animals over weeks or months, continuous performance monitoring, and gradual adjustment of task parameters throughout the training^9–11^. Additionally, training protocols vary widely between laboratories and often depend on subjective experimenter criteria, limiting their reproducibility^12^. To confront this experimental bottleneck, high-throughput multi-box systems mitigate these constraints by scaling up and automating training procedures, enabling the parallel testing of multiple animals^13–16^. Yet, these multi-box systems still rely on human intervention to transfer animals from home cages to operant boxes, involve substantial building and maintenance costs, and require considerable space. Such animal handling, apart from being time-consuming, can affect animals’ internal states by interrupting ongoing home cage activity^17,18^, constraining their training schedules, and in some cases, abruptly changing the context (e.g., training rooms separated from housing rooms), potentially impacting performance and learning^19^. Additionally, to optimize the time invested in training, animals are commonly food or water-deprived for prolonged periods, which can cause marked distress along with physiological and behavioral changes^20,21^. Lastly, observing animals only under these highly stereotyped conditions (e.g., once daily, at the same time, with similar motivational state) may overlook important contextual information, such as fluctuations in motivational level, circadian variables, and the circumstances under which they choose to engage in training^22^.

To overcome some of these limitations, several laboratories are shifting towards fully automated home cage training systems that minimize human intervention by making animals train autonomously^23–40^. Some of these systems incorporate continuous monitoring of home cage activity and prioritize the study of social interactions and ethological behaviors^32,38^, but rely on simpler behavioral tasks with limited translational relevance. Other approaches focus primarily on operant task training but either rely on single-housed animals^25,26,31,34,37,39,40^, or implement tasks directly inside the home cage, which limits the type of tasks that can be performed and the experimental control (e.g., reduced control over task access and increased distraction from conspecifics)^23,27,28,30,36^. Finally, a few systems separate group-housed home cages from the operant box, allowing finer control over task execution while preserving social housing, but they either use commercial software solutions^29,35,38^, or offer limited software development^24^. Additionally, none of these systems is modularly designed to support integration across multiple behavioral paradigms, diverse operant box designs, and different acquisition systems. Despite substantial progress, no existing platform simultaneously combines i) continuous autonomous training in complex cognitive tasks, ii) modularity and compatibility with different operant paradigms and acquisition systems, iii) group housing with preserved social interactions, iv) longitudinal monitoring of spontaneous home cage behavior, v) remote user-friendly access for close-up supervision, and vi) affordability (Supplementary Table 1).

Here, we developed the Training Village (TV), an open-source system designed with all these features for the continuous automated training of both mice and rats in almost any freely-moving task. In the TV, a group of animals lives freely in enriched arenas displaying complex social interactions while individually accessing the operant box to perform cognitively demanding tasks at any time. Given its modularity, the TV is designed to be *wrapped around* existing behavioral systems without requiring users to change their current task paradigms. The system organizes the training and testing of each animal by continuously monitoring task performance and automatically adjusting parameters without direct human intervention. Moreover, it provides remote supervision of animals’ behavior, alarm system, and allows manual intervention when needed. Extensive testing showed that animal groups share the use of a single operant box very efficiently and uniformly, and engage in diverse cognitive tasks with high and stable performance over months. In sum, the Training Village has the potential to transform rodent cognitive training into a scalable and efficient process. By enabling longitudinal testing of the same individuals over extended periods, it provides a powerful framework to study the mechanisms of cognitive decline. Moreover, the integration of home cage activity and behavioral testing allows the investigation of individual differences in behavior. Together, these features have the potential to accelerate the urgent search for new behavioral markers of brain function and dysfunction.

## Results

### An automated training system with continuous data acquisition

To enable continuous and fully automated training of rodents in cognitively demanding tasks, we developed the Training Village (TV), an open-source system that autonomously trains and tests rodents in cognitive tasks while simultaneously monitoring their home cage activity. The TV is modular and composed of three main elements: The home cages, the operant box, and a corridor connecting the two. The *home cages,* mimicking the Eco-HAB design^41^, typically consist of several home cages connected by tubes, where groups of animals live in enriched environments that emulate their natural burrow habitat. Each animal is implanted with a radio-frequency identification (RFID) microtransponder for individual identification, and RFID sensors are installed in the connecting tubes, enabling tracking of the cage in which each animal is located at any point in time. The *operant box*, where animals are trained and tested, runs a specific trial-based behavioral paradigm. Given its modularity, the TV is compatible with multiple types of operant chambers, allowing a broad range of task designs. We have validated two operant box designs: the standard three-port box commonly used in two-alternative decision-making tasks^42–47^, and a T-shaped maze with a touchscreen on one end and a reward port on the other end^48^. Lastly, the *corridor,* which connects the home cages to the operant box through a transparent tube equipped with two motorized doors and a double identification system based on an RFID reader continuous video analysis of the animals’ activity inside the corridor (Fig. 1a, Supplementary Fig. S1, and Supplementary Video 1). The corridor plays a critical role in the system by ensuring animals enter the operant box one at a time. Importantly, animals do not need to be completely water-restricted. They can have free access to slightly sour water containing a small amount of citric acid (CA) in their home cages, and are rewarded with sweetened water in the operant box. This approach provides sufficient task motivation without strict water restriction^49^ and ensures that animals maintain healthy weights.

**Figure 1:**
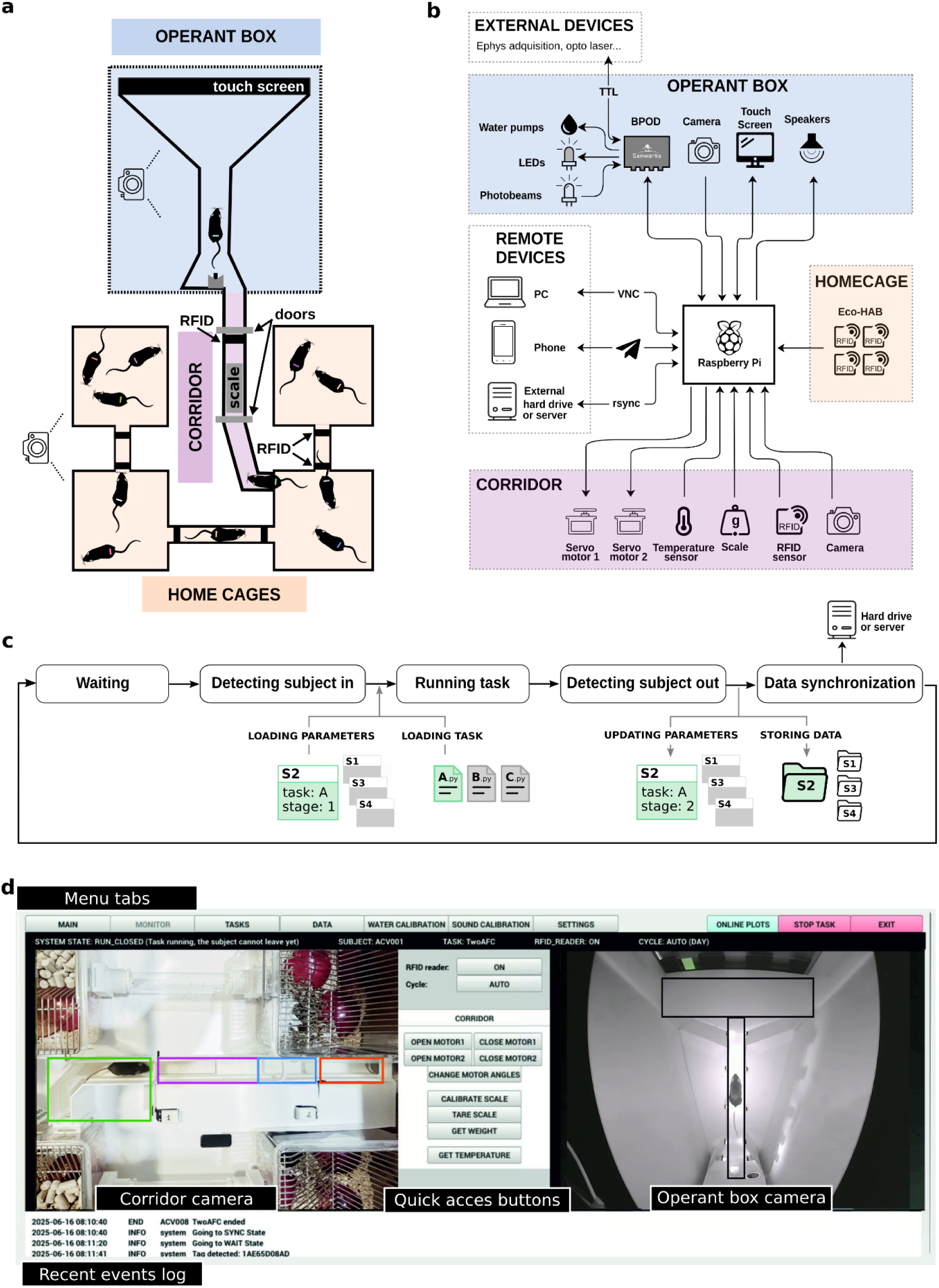
Training Village hardware and software. **a**, Schematic layout of the TV, composed of three main elements: the home cages, the operant box, and the corridor that regulates access to the operant box. Cameras are placed above the system. **b**, Diagram of hardware connections. The TV is controlled by a Raspberry Pi, which sends and receives signals from electronic devices in the corridor and operant box, and controls the remote devices. **c**, Diagram of the main states of the system and data flow. When an animal is detected in the corridor, the corresponding parameters are loaded, and the behavioral session is initiated. After task completion, the generated data is automatically added to a global DataFrame that contains all information about the subject. This information is used to update a parameter dictionary for that subject, which adapts task difficulty and training stage based on the subject’s performance. Finally, data is backed up to an external device. **d**, Graphical User Interface of the TV. Shortcut buttons allow direct system control, while multiple tabs provide intuitive interaction and user-friendly access to the system settings and data, and allow the manual initiation of behavioral tasks. Two cameras continuously monitor activity in the operant box and corridor. Real-time behavioral plots and event logs are also available to inform the user about the system status quickly (Supplementary Fig. S3).

The TV’s central control unit is a Raspberry Pi 5, which continuously runs a custom-made open-source software. The Raspberry Pi is a robust, low-cost (∼100€), low-power (∼15 W), and compact computer. Its standardized architecture guarantees reliable and seamless integration of software and hardware components while offering great scalability when building multiple systems. The Raspberry Pi controls the corridor doors, reads data from RFID and other sensors, records video from two cameras, manages the overall system workflow, communicates with the task-control devices, and handles both data storage and remote communication (Fig. 1b). Because the system is modular, task control can be implemented in multiple ways. We used Bpod (Sanworks), an open-source behavioral control system that offers sub-millisecond timing precision, a convenient state-machine framework for designing tasks, and automatic real-time event logging to CSV during each trial. Other controllers, such as Arduino, pyControl^14^, or Harp (Open Ephys), can be adapted as long as they meet equivalent data-logging standards. For tasks that do not require high temporal precision and can be driven directly from the Raspberry Pi’s GPIO, no external controller is required; the Raspberry Pi can execute the entire task logic.

The TV implements a state machine for the control of the access and exit to the operant box and to run task variants corresponding to different training stages. In the initial state, the system waits for an animal to enter the corridor and cross the door one. Once a subject RFID tag is detected near the door two and the camera above the corridor does not detect any other animals in the corridor, the system consults a local database to retrieve subject’s assigned task parameters and checks whether enough time has elapsed since the last training session. If this condition is met, door one closes and the door two opens, allowing the animal to access the operant chamber, which triggers the door two to close. The TV automatically starts the behavioral task, and after a configurable minimum session duration, the door two opens and the animal is free to exit (Supplementary Fig. S2a). On exit, before reaching door one, the animal steps on a scale which records its weight and triggers the door one to open and the door two to close, confirming the animal’s return to the home cage. Finally, the system returns to the initial state (Fig. 1c and Supplementary Fig. S2a).

After each session, task data (events, state transitions, timestamps, etc.) is automatically saved in a CSV file. A customizable Python script processes this data to update the task parameters for the next session and stores them in a JSON file. Task CSV and video data are uploaded to a remote server or external hard drive (Fig. 1c). The TV automatically monitors the evolution of task performance across sessions and advances each subject through training stages according to criteria set *a priori* (e.g., sessions’ accuracy, number of trials completed, etc.). This standardization of the training criteria enables dynamic task progression tailored to each subject’s performance, entirely without experimenter intervention. This level of automatization minimizes human biases, common in manual training, and ensures reproducible conditions for investigating learning across different groups of mice. The system also provides the option of changing parameters manually in a flexible manner.

Given its 24/7 operation, the TV is designed to be predominantly operated remotely. Access to the Raspberry Pi controlling the TV can be established via any remote desktop software (e.g., AnyDesk) or through VNC. The graphical user interface (GUI) enables user-friendly monitoring and control of both elementary and advanced system functions (Fig. 1d, Supplementary Fig. S3). First, the user can visualize the state of the system via two cameras, one imaging the corridor and the home cages, and another imaging the operant box. Second, the GUI includes online plotting during behavioral sessions, across-session summary reports monitoring training progress, and weight variation and temperature reports to monitor animal health (Supplementary Fig. S3e). All these plots are easily customizable. Third, the GUI allows for easy and organized access to the videos of previous sessions to quickly identify technical issues or undesired behavior. Finally, the system features an alarm system via messaging apps (e.g., Telegram), allowing users to receive real-time notifications on their mobile devices and observe and control the system from any remote location (Supplementary Table 2). This infrastructure enables efficient management without constant physical presence.

The entire system is open-source and has been designed in a modular and flexible way to allow an easy implementation of user-defined behavioral tasks and training progress settings. The materials list, 3D designs, software, updated and extensive documentation are available through the TV website. The cost of the TV is approximately 5500€ (1000€ for the corridor; 1500€ for the home cage, including the Eco-HAB reader; 2500€ for the basic operant box, including the Bpod; and 500€ for the touchscreen), which is more affordable than commercial automated training systems^26,28,50^. Assembly of the entire system can vary between 2 and 7 days, depending on the experience of the installer. We have tested the TV for both mice and rats with scaled versions of the corridor and operant box to accommodate each species’ requirements (Supplementary Fig. S1, and Supplementary Video 1).

### Efficient and balanced access to the operant box

The Training Village is a convenient way to train groups of rodents in complex tasks that take many sessions to be learned. Nevertheless, it also presents several challenges as training is individually self-paced (an animal only trains when it decides to enter the operant box), and a single operant box must be shared by the entire group. This could *a priori* generate several inefficiencies, such as access bottlenecks when multiple animals attempt to enter simultaneously, or periods of underuse when all animals are resting. We next show that by adjusting the size of the animal group and constraining the timing and length of the sessions, the animals self-organize and use the system in a highly efficient and systematic manner, generating high-quality behavioral data at a fast rate.

We tested a total of 12 groups in the TV (11 mouse groups and 1 rat group, comprising a total of 121 subjects). Every animal performed multiple entries into the operant box distributed throughout the day (see example group in Fig. 2a). In addition, animals used the operant box intensively. For example, Fig. 2b, shows the time-resolved operant box occupancy for a group of 12 mice, which fluctuated around 80% of the total time from the start of training. As expected, the average operant box occupancy increased with group size, but this increase was marginal for groups above eleven animals (Fig. 2c). As the TV can work all days of the week, an occupancy of around 80% could potentially yield an average of ∼134 hours/week of behavioral task data.

**Figure 2:**
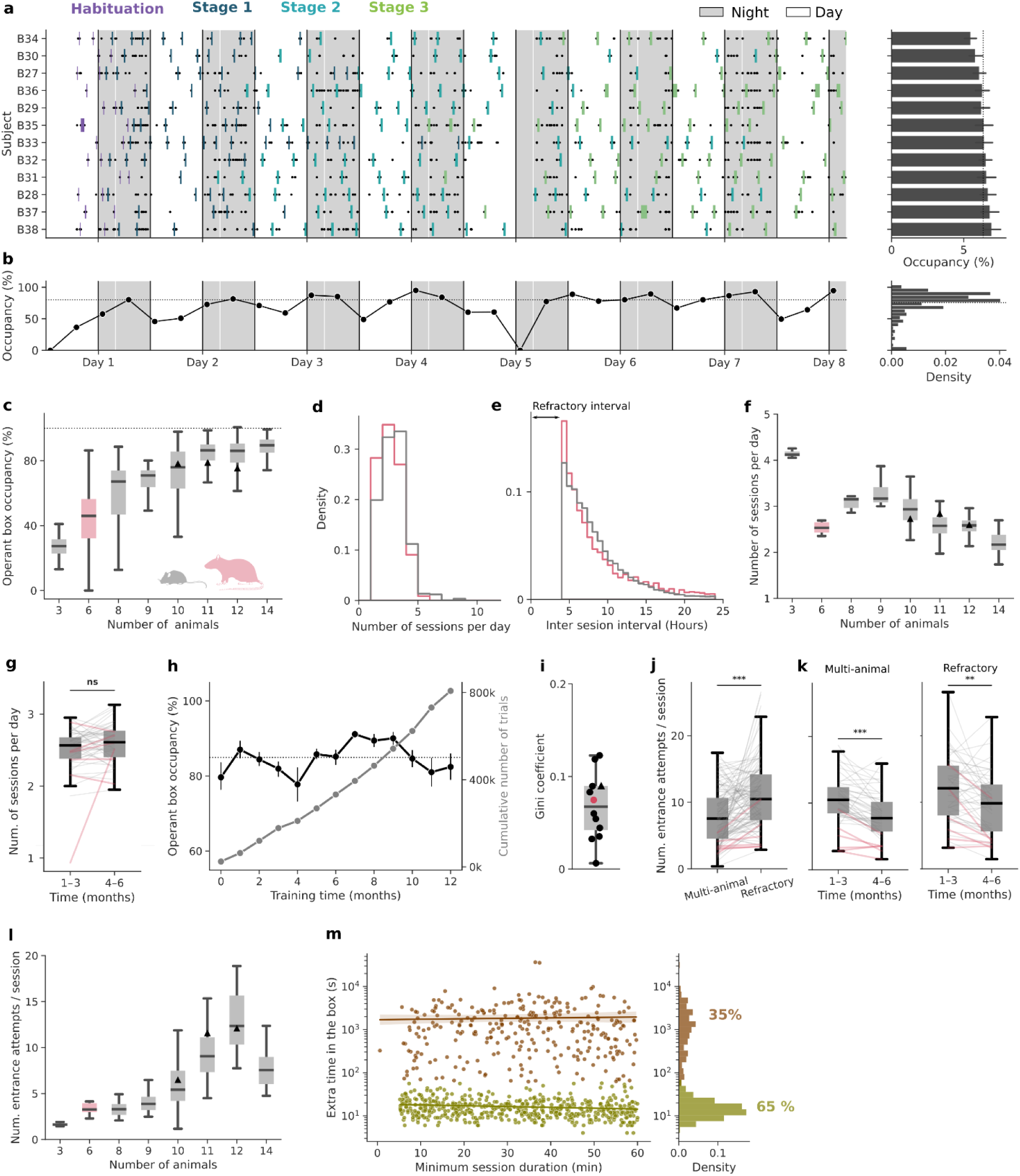
Self-regulated behavior in the TV results in an efficient and well-distributed use of the operant box. **a,** *Left*: Raster plot of entries into the operant box (rectangles, colors represent training stages) and entrance attempts (black dots) during the first 8 days of training (Group 6, n=12). Gray areas represent nighttime. *Right*: Histogram of individual occupancy of the operant box by the same mouse group shown in a (same y-axis). Each bar represents the mean individual occupancy of the box across the entire experiment (error bars indicate the 95% CI). The self-organized distribution of the box usage time is very close to a uniform distribution (dashed line). **b,** Evolution of the operant box total occupancy during the same period shown in a (left), and its histogram over the entire experiment (right). Connected dots indicate the mean occupancy in 6-hour bins. The drop on day 5 was due to system maintenance. The dashed line represents mean occupancy over the entire experiment. **c,** Operant box average occupancy as a function of group size (sizes 3 to 9 correspond to single groups, while sizes 10–14 aggregate multiple groups). Rat cohort indicated in pink. The black triangles represent the example group shown in panels a-b, whose size varied across time. **d,** Histogram of the number of daily sessions per animal computed separately for mice (gray) and rats (pink). **e,** Histogram of the inter-session interval (i.e., time between successive behavioral box entrances of the same subject). No entrances occurred within the first 4 hours after a session ended due to the imposed refractory interval. **f,** Average number of daily sessions as a function of the group size. **g,** Average number of daily sessions over time (grouped by trimesters). Data is from groups 4, 5, 6, 7, and 9, which ran on the TV for at least 6 months. Paired t-test: *t* = -1.86, *p* = 0.07. **h,** Monthly average occupancy (black) and cumulative number of task trials (gray) versus experiment time (error bars indicate 95% CI; Group 7). **i,** Quantification of the distribution of box usage across animals by the Gini coefficient for the different groups tested in the TV. **j,** Number of failed entrance attempts per session (i.e., per successful entrance) calculated by type of attempt. Paired t-test t = -5.7, p < 0.001. **k,** Failed entrance attempts per session calculated separately during the first and second trimester of data collection. Same dataset as in g. Paired t-test: *t* = 5.6, *p* < 0.001 (left); *t* = 3.3, *p* = 0.002 (right). **l,** Number of failed entrance attempts per session shown as a function of the group size. Only *multi-animal* attempts were considered for this analysis. **m,** Time spent inside the operant box after the door opened as a function of the door opening time (left) and its histogram (right). Sessions were classified as fast exits (green, extra time <1 min) or slow exits (brown, >1 min extra time). Each dot represents one session. Regression lines for fast (slope = -0.07, *p* = 0.002) and slow exits (slope = 4.4, p = 0.73). Analysis is based on Groups 5 and 6 (n=829 sessions). Boxes show the median and IQR across groups (panel i), days (c), or subjects (f, j, k, l).

In order to regularize the duration of the sessions and to prevent animals from entering the box without sufficient motivation (i.e. thirst), we imposed (1) a minimal session duration during which the door remained closed and subjects had to stay in the operant box; (2) a minimum inter-session interval, called the *refractory interval* (typically 4 hours), during which animals could not re-enter the operant box (Methods). With these constraints, on average, individual animals accessed the operant chamber several times per day (between 1 and 5 sessions, mean = 2.5 for mice and 2.2 for rats; Fig. 2d). The inter-session interval exhibited the shape of a shifted-exponential distribution with a median of 7 hours (95.5th percentile = 16.9 and 19.2 for mice and rats, respectively; Fig. 2e). The daily number of sessions per subject increased for smaller groups as the competition to enter the box diminished, except for rats which, compared to groups of mice of similar size, completed fewer daily sessions (Fig. 2f). Thus, in general, animals preferred to distribute their daily work in the task across several shorter sessions, rather than completing a longer single session as is commonly done in standard manual training. We then tested the effects of removing the time constraints (minimum session duration and the refractory interval), so that subjects could come into the box whenever it was empty and leave it anytime (Group 12, n = 10 mice). Interestingly, without restrictions, mice increased the average number of daily sessions but performed fewer trials per session, ultimately resulting in a similar number of daily trials (Supplementary Fig. S4a). Additionally, the variability across subjects increased for both the box occupancy and trials per session (Fig. S4b-c). Hence, the time constraints worked as a regularizer of the number of these two variables, but did not affect the total number of daily trials. Next, we asked whether this high motivation to enter the box was maintained throughout long experiments. We found that the number of daily sessions did not vary significantly over months (Fig. 2g) and that the system could generate thousands of task trials at a steady rate over long periods, making it particularly well-suited for long longitudinal experiments (Fig. 2h).

Given that mice and rats are nocturnal, we examined the circadian modulation of operant box usage and found that during both the light and dark cycles, animals showed high occupancy with a slight but significant increase during the night (Supplementary Fig. S5a-b). The number of sessions performed and the number of entrance attempts per session were also significantly higher during the night, although both remained substantial during the day (Supplementary Fig. S5d-f). Elevated daytime activity could, in principle, result from nighttime operant box overcrowding, forcing animals to access the box during the day. To test this possibility, we quantified day–night occupancy differences as a function of group size. Occupancy remained comparable between light and dark phases across all group sizes tested (Supplementary Fig. S5c), indicating that daytime operant box usage was not driven by limited nighttime access, but instead reflects sustained engagement with the task throughout the circadian cycle.

Mice and rats form hierarchies in which a few individuals can exert dominance and territoriality over the more submissive conspecifics^51–53^. This innate behavior could lead to a few individuals dominating access to the operant box, which is a limited resource in the environment. However, the individual occupancy of the box was almost uniformly distributed across animals (Fig. 2a, right). Regardless of the group size, all groups distributed the use of the box in an egalitarian manner, as reflected by low Gini coefficients (all below 0.15, where 0 indicates perfect equality and 1 maximal inequality) (Fig. 2i). This uniformity was likely facilitated by the imposed refractory interval between sessions, which forced animals to space apart their entrances. Interestingly, after removing this restriction in an already trained group, the occupancy distribution remained broadly uniform, although it became significantly less egalitarian (Gini = 0.03 vs 0.07, p < 0.001; Supplementary Fig. S4b). This suggests that, while access restrictions promote a more balanced usage, animals trained under these constraints do not develop strong dominance patterns in operant box access once the restriction is lifted.

Although all animals used the operant box regularly, accessing it required substantial persistence. Each subject made an average of 23 failed entrance attempts per day (approximately 9 attempts per successful entrance). Entrance attempts occurred when an animal entered the corridor (i.e., the box was empty) and was identified by the RFID reader, but it could not access the box because (1) either several animals tried to enter simultaneously in the operant box (*multi-animal* attempts) or (2) because the attempt occurred during the refractory interval (*refractory* attempts), being the latter more predominant (*p* < 0.001; Fig. 2j). Over time, animals learned to coordinate their entrances with cage mates, as shown by the significant reduction in the daily *multi-animal* attempts number(*p* < 0.001; Fig. 2k left), as well as the refractory interval rule, significantly reducing attempts during this period (*p* = 0.002; Fig. 2k right). As expected, larger groups showed in general a larger number of entrance attempts as the competition to access the box increased (Fig. 2l). Altogether, we concluded that group sizes around 10 subjects provided a good balance between maximizing the operant box usage (Fig. 2c) and the number of daily training sessions (Fig. 2f), while keeping the number of failed entrance attempts low (Fig. 2l).

We observed that mice were highly motivated and persistent in accessing the operant box. However, once inside, it remained unclear what triggered their decision to exit. In two groups of mice (Groups 5-6, n= 21), we randomly varied the minimum session duration, which was determined by the door opening time (uniform distribution between 5 and 60 minutes). Under these conditions, once the door opened (the motor generates an audible cue), animals could decide to leave immediately and end the session, or to remain inside and continue training. In most cases (65% of the sessions), mice exited immediately after the door opened (< 1 minute, Fig. 2m right), indicating that door opening was a strong cue to trigger their exit. These rapid exits occurred regardless of how long the mouse had been training, but tended to be slightly faster in longer sessions (Fig. 2m left, slope = -0.07, *p* = 0.002). In the remaining 35% of sessions, animals stayed for longer and more variable periods after the door opened. These extended stays were slightly more frequent in sessions with longer door opening times (32% vs 37% for shorter vs longer duration relative to the mean; *p* < 0.001), yet their duration was independent of the door opening time (slope = 4.4, *p* = 0.73), suggesting they were likely due to animals missing the door opening cue, rather than a deliberate choice to continue training. In total, the Training Village takes advantage of mice’s natural curiosity and preference for exploration, which drives them to try to enter the box continuously. We found that even when they had unrestricted availability of plain water in their home cages and the access to the box was discontinued, animals persisted attempting to enter it after several weeks (Supplementary Fig. S5g). Fortunately, this natural instinct to explore was accompanied by a tendency to return to their home cage soon after entering the box. This prevents the undesirable scenario in which mice stay much beyond the time of task engagement (blocking other mice from using the box) and allows control to a large extent of the box occupancy by timing when the door opens.

### Animals learn and engage in multiple cognitive tasks using the Training Village

Having shown that using the TV, animals accessed the operant box continuously, we next asked to what extent they engaged in the task once inside the box. Because animals self-paced their entrances at variable times during the day, we wondered if this variability would result in heterogeneous levels of engagement across animals or sessions. To examine this, we trained distinct groups of animals in three behavioral paradigms with different cognitive demands.

First, mice and rats were trained in a visuospatial delayed-response three-alternative choice task (3AFC) during which they briefly viewed a stimulus displayed on a touchscreen and had to report the stimulus location by nose-poking the screen at the corresponding position (Fig. 3a, Groups 1-8 with n= 92 mice; Group 9 with n=6 rats). Stimulus duration varied from trial to trial, producing a variable mnemonic delay. Correct responses were rewarded at the end of the maze opposite the touchscreen (Fig. 3a). Second, we also trained mice in a visual discrimination two-alternative forced choice task (2AFC) in which, after nose-poking in a central port, the two adjacent ports were illuminated at different light intensities and mice had to poke in the brighter port to receive the reward (Fig. 3b; Group 12, n=10). An auditory frequency discrimination task, named the cloud of tones task^54,55^, was also tested with the same group of animals. Finally, we trained a group of mice (Group 11, n=10) in a two-armed bandit task (2AB) in which mice had to choose between left and right ports that delivered rewards with complementary probabilities, which reversed periodically across blocks (Fig. 3c)^47^. In all tasks, animals received as a reward a drop of sweetened water for every correct response. Due to the nature of each task, the mean number of daily trials performed varied greatly: 180 trials in mice performing the 3AFC, 666 in 2AFC, and 690 in 2AB. It also depended on the species, with mice performing more trials than rats (108 trials in 3AFC) (Supplementary Fig. S6a).

**Figure 3:**
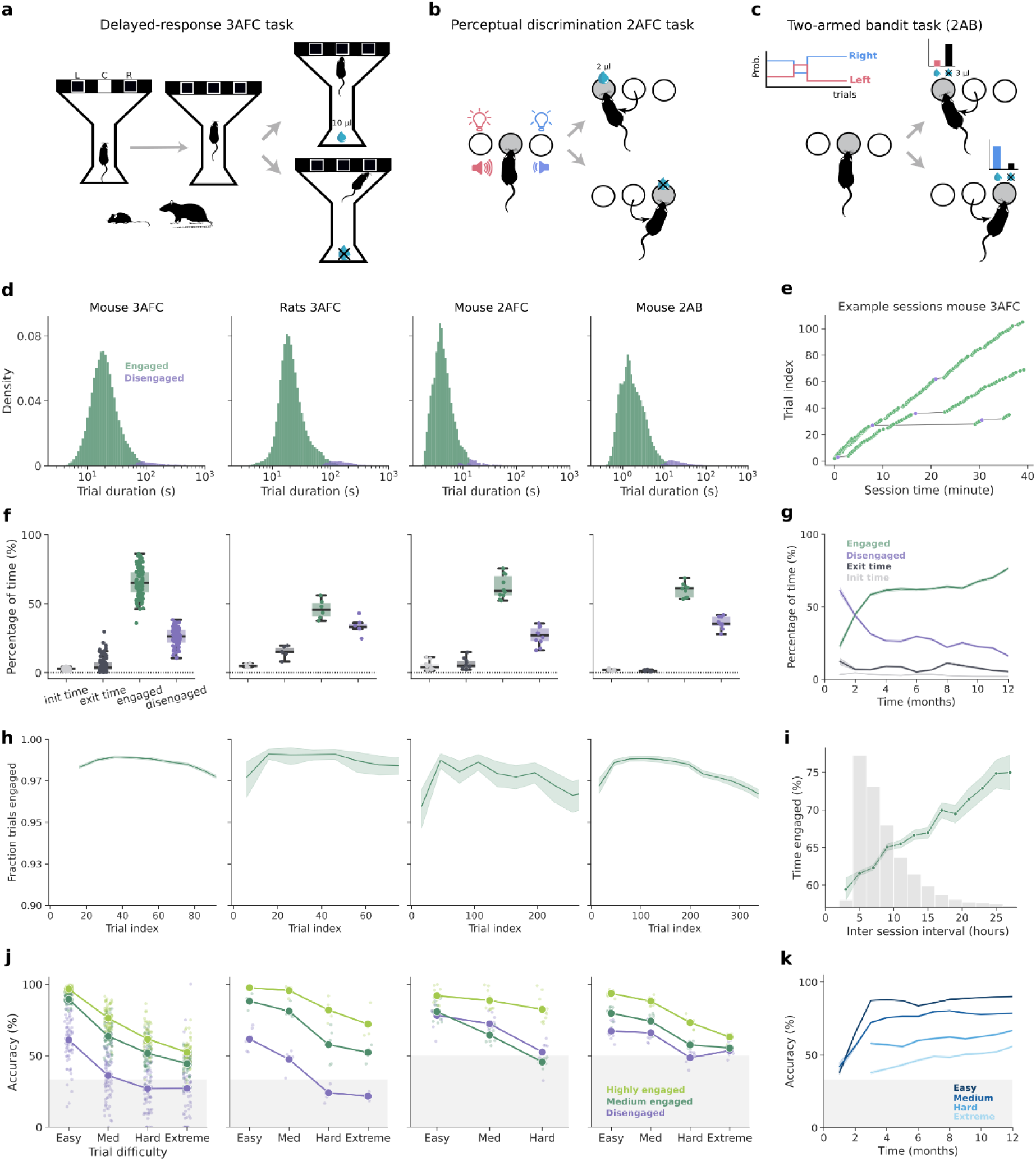
Animals are consistently engaged in different cognitive tasks tested in the Training Village. **a-c,** Schematics of the behavioral tasks implemented and tested in the TV. **a,** *Visuospatial three-alternative choice delayed-response task* (3AFC), adapted for mice and rats. **b,** *Two-alternative discrimination task* for mice (2AFC). **c,** *Two-armed bandit task* for mice (2AB). Panel columns in d, f, h and j correspond to each task; see panel titles. **d,** Distributions of the trial duration obtained from all animals in each task. 3AFC Median = 19.1 s, Groups 1-8, n= 88 mice, N= 10^5^ trials; 3AFC Median = 20.6 s, Group 9, n= 6 rats, N=84,670 trials; 2AFC Median = 4.1 s, Group 12, n=10 mice, N = 49,727 trials; 2AB Median = 1.65 s, n = 9 mice, N = 450,382 trials. In the 2AB task, we excluded a portion of the trial to avoid confounds, as a consequence, trial durations appear shorter (see Methods for details). **e,** Trial index as a function of trial onset in three example sessions from the same mouse carrying out the 3AFC task. Notice that trials containing long pauses are classified as disengaged (purple). **f,** Averaged percentage of time spent by each subject in each of the four engagement states: time to complete the first trial, time from door opening to the animal’s exit, time engaged and time disengaged. Dots represent subjects, and boxes show the IQR with the median. **g,** Percentage of time spent in the four engagement states (defined in f) across months (only Group 7, n = 12). Lines represent the monthly averages and shaded bands indicate the SEM. **h,** Fraction of engaged trials as a function of trial index. Lines represent averages across bins (10-trial bins in 3AFC and 30-trial bins in 2AFC and 2AB). **i,** Fraction of engaged trials over a session as a function of the preceding inter-session interval (ISI). Dots connected by a line represent the mean fraction across sessions with similar ISI and shaded bands the SEM. Histogram of ISIs is shown in gray. **j,** Choice accuracy as a function of trial difficulty and different levels of engagement. More details of the engagement calculation and trial difficulty categorization are in Methods. Small dots represent individual subjects. Larger dots connected by lines represent mean ± SEM. The gray area represents chance-level accuracy. **k**, Choice accuracy across months split by trial difficulty (Group 7, n = 12 as in panel g). Lines represent monthly averages, and shaded bands indicate the SEM. Equivalent plots for the auditory 2AFC task are shown in Supplementary Fig. S6f (corresponding to panels d, f, and j).

As animals self-initiated the trials in all tasks, we quantified their task engagement based on the trial duration, which included the time they initiate and complete each trial. For every subject, we classified trials as *disengaged* when their duration exceeded two standard deviations above the mean (Fig. 3d), and *engaged* otherwise. Although the mean trial duration differed substantially across tasks, in all cases subjects exhibited alternating epochs of engagement, characterized by sequences of consecutive trials carried out at fast speed, and periods of disengagement when they paused for a variable interval (Fig. 3e, Supplementary Fig. S6b). On average, only a small fraction of trials per session were disengaged (notice the long tail in the distribution of trial durations in Fig. 3d). Although few in number, disengaged trials could account for a large portion of the time spent in the operant box. To finely assess how efficiently animals used their time inside the box, we aggregated the duration of engaged and disengaged trials into engaged and disengaged time, respectively. Additionally, we computed the latency to complete the first trial after entering the box, and the latency to exit once the door opened (minimum session time elapsed). Across tasks, on average, animals spent ∼64% of the time engaged in the task (3AFC: 65%; 2AFC: 62%; 2AB: 60%) (Fig. 3f), and ∼28% of the time disengaged, while the remaining time was spent completing the first trial (3%), and exiting the operant box (5%). Rats were generally less engaged (46%) than mice, and showed longer exit latencies (15%, Fig. 3f). As expected, we observed that higher percentages of time engaged in the task led to a larger number of trials (Supplementary Fig. S6c). We also observed that animals spent more time engaged in the task during the night sessions, except for mice trained in the 3AFC task (Supplementary Fig. S6e). Notably, in animals trained over long periods (Group 7, n= 14), task engagement gradually increased over a year, suggesting that extensive experience leads to more sustained engagement (Fig. 3g).

The slow drift of behavioral variables over the course of the session is commonly observed in rodent experimentation. For example, task engagement typically decreases with time, reflecting a shift in their motivational state as animals transition from high deprivation to increasing satiation^56–58^. In the TV, we found that the fraction of engaged trials changed slightly during the session and followed a consistent pattern across tasks: it started low, reached the peak after a few trials, and then gradually decreased throughout the session (Fig. 3h). A similar trend was observed when using trial duration as a proxy for task engagement (Supplementary Fig. S6d). Overall, this pattern suggests that animals begin the session slightly distracted, probably exploring the box, then quickly focus on the task, reaching peak engagement with the fastest reaction times, which then declines slightly over time, possibly due to satiation. Engagement also showed a systematic variation across sessions dependent on the length of the inter-session interval: as expected, animals were more engaged in sessions following longer inter-session intervals (Fig. 3i). We also found that engagement correlated with task performance. For each task, we plotted psychometric curves showing how choice accuracy decreased with the difficulty of the trial (Fig. 3j): in the 3AFC task, accuracy decreased with longer mnemonic delays; in the 2AFC task, it decreased with smaller intensity differences; and in the 2AB task, it decreased with the similarity of the reward probabilities between ports. Notably, engagement was associated with higher choice accuracy in all tasks (Fig. 3j). Finally, we found that engagement and mean accuracy continued increasing over long training periods (Fig. 3g,k), demonstrating the platform’s suitability for complex tasks that require long training and testing periods.

Finally, we compared task performance in mice trained in the 3AFC task either automatically using the TV or manually in one-hour daily sessions. Despite distributing the daily work across several sessions (Fig. 2d), the total number of trials per day in the TV was, on average, similar to manual training (medians: TV = 182 trials, manual = 167 trials; Supplementary Fig. S7a-b). Furthermore, self-trained animals showed slightly higher average accuracy (68%) than the manually trained mice (64%, p = 0.02, Supplementary Fig. S7c).

### Linking home cage behaviors with task activity

Using the Training Village we could monitor animals’ activity, not only in the operant box while performing the task, but also in the home cages by integrating the system with the Eco-HAB^41^. In one TV setup for mice, we connected four cages with tunnels equipped with RFID antennas, enabling continuous tracking of the cage each animal occupied at any given time. To infer different types of home cage behaviors, we placed distinct elements on each of the four cages: (1) *eating* behavior was inferred from the presence of an animal in the cage containing food and citric-acid-flavored water; (2) *sleeping* behavior from the cage that contained nesting material and shelters; (3) *playing* behavior from the cage that offered running wheels, tubes, and wooden sticks for enrichment; and (4) *task seeking* behavior from the cage that was empty and provided access to the operant box (Fig. 4a). Continuous activity monitoring revealed that, between task sessions in the operant box, mice explored all compartments multiple times per day (Fig. 4b), although they spent the majority of the time in the sleeping cage (Fig. 4a, bottom), especially during the light phase (Fig. 4b). Circadian modulation of home cage activity was quantified by the average number of corridor crossings per hour, confirming higher locomotor activity during the dark phase (Fig. 4c-d, p < 0.001). Notably, despite this circadian modulation, mice remained relatively active during the light phase (Fig. 4c), consistent with their continued access to the operant box during this period (Supplementary Fig. S5). We next examined whether task execution altered the natural circadian rhythm of mice. To test this, we compared corridor activity under standard TV conditions (i.e. with an active task) to a no-task condition (i.e. with no access to the operant box and *ad libitum* water in the home cage). The circadian activity pattern was largely preserved across conditions (17.8 vs. 17.1 mean number of cage changes per day; *p* = 0.06; Fig. 4c-d). Although task availability was associated with a small reduction in overall home cage activity, the effect size was small (17.8 vs. 17.1 mean number of cage changes per day; *p* = 0.03; Fig. 4d), indicating that task performance does not seem to introduce major sleep-cycle disruptions that could raise concerns about the animals’ overall welfare.

**Figure 4:**
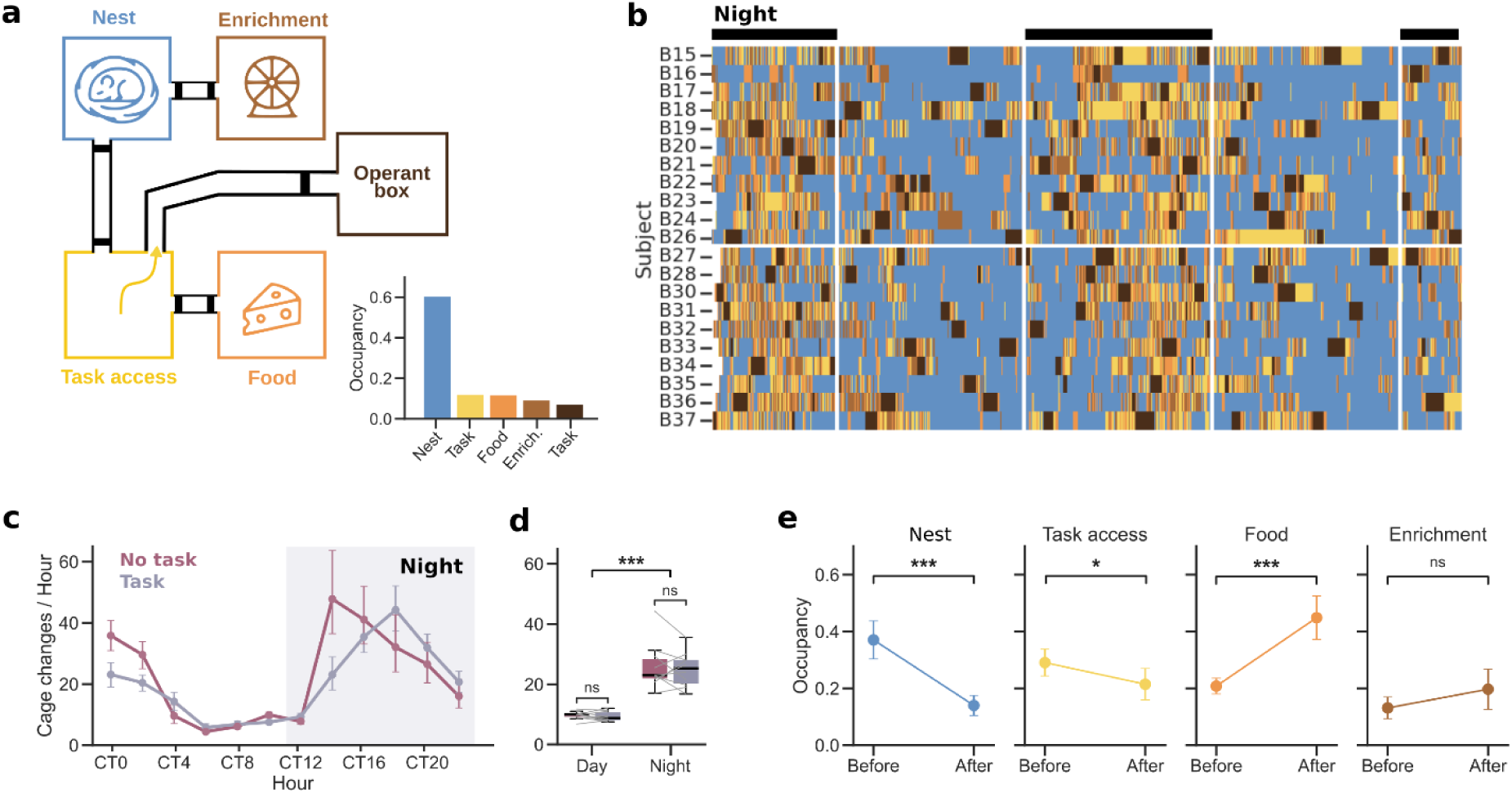
Home cage behaviors are modulated by task activity. **a,** Schematic of the TV + Eco-HAB spatial arrangement. Bottom right: Averaged daily occupancy for each box. **b**, Raster plot showing individual mouse location across the four home cages and the operant box over two days in two different TVs (Group 5 n = 11, and Group 6 n = 10, separated by the horizontal line). Each row represents one mouse, and colored boxes represent periods during which a mouse remained in one of the different types of cages described in panel a. Black bars on top indicate night phases (limited by vertical lines). **c,** Home cage activity quantified as the median number of cage changes per hour plotted in circadian time. Data is split by system status (task: normal access to the operant box; no task: no access to the operant box and plain water *ad libitum* in the home cages). Dots represent the mean across subjects; error bars represent 95% CI. The gray band represents the nighttime period. **d,** Median number of cage changes during day and night phases, split by system status. Lines represent individual subjects; boxplots show the median and IQR. Effects of task status, light cycle, and their interaction were assessed using a linear mixed-effects model with subject as a random intercept: task status *β* = -4.7, *p* = 0.06; cycle *β* = 12.6, *p* < 0.001; interaction *β =* 2.2, *p* = 0.6. Significance markers indicate results from paired t-tests. **e,** Cage occupancy during the 5 minutes before and after each behavioral session. Dots represent the mean across subjects, and error bars show 95% CI. Paired t-test: *t* = 8.9, *p* < 0.001 (Nest), *t* = 3.3, *p* = 0.04 (Task access), *t* = -8.4, *p* < 0.001 (Food); *t* = -1.7, *p* = 0.5 (Enrichment). All *p*-values were corrected for multiple comparisons using Bonferroni. Panels c-e show data from Group 5, n=11.

We next examined home cage occupancy preferences immediately before and after task sessions (5-minute periods). We found that mice tended to be most often in the sleep cage before the task (*p* < 0.001), and in the eating cage after (*p* < 0.001) (Fig. 4e). The occupancy of the cage providing access to the operant box was slightly higher before than after (*p* = 0.04), probably due to their entrance attempts, but remained constant for the enrichment cage (*p* = 0.5). These results suggest that when mice are sleeping, they may wake up by the sound of the door opening or by the arrival of another animal that had just finished training, which provokes them to try entering the operant box. After training, having drunk sufficient water, they eat. Together, our results illustrate the potential of TV to examine how behavioral tasks influence home cage activity and to understand the impact that contextual variables have on the task, ultimately providing a more comprehensive view of the relationships between individual cognition, learning, and innate behaviors.

## Discussion

Training rodents in complex cognitive tasks with high translational relevance remains a major experimental bottleneck in neuroscience. Here, we introduced the Training Village as a powerful system that takes rodent training to a new level of automation. It enables stress-free, 24-hour autonomous training without human intervention while animals live and engage socially in their home cages. In comparison to other systems^23–32,34–40^, the TV stands out due to its modularity and compatibility across a variety of tasks and to its effort in closely monitoring the wellbeing of mice both in the home cage and during the task.

The most common solution to accelerate rodent training in complex tasks is to build scaled-up behavioral racks containing *N* behavioral boxes, enabling the parallel training of *N* animals^13–15^. Despite the clear benefits, such systems present several limitations: First, each experimenter can only run a finite number of operant boxes (*N_max_*) simultaneously, constrained by the manual workload required between sessions (e.g., remove animal from box, bring new one, weight it, annotate, identify protocol and run), such that *N_max_* = session duration/manual workload duration. Second, when managing dozens of animals across multiple protocols and boxes, human errors inevitably occur. These errors can be difficult to trace back, potentially leading to increased variability in the task learning success or wrong conclusions. Third, equipment costs scale linearly with the number of boxes. Fourth, each operant box necessarily imposes a maintenance cost which includes cleaning, calibration, and routine repairs. Identifying problems in one out of *N* boxes is challenging, often resulting in less finely controlled instrumentation which increases the sources of variability and limits reproducibility. Fifth, scaled-up systems can occupy substantial physical space, a limited resource in many animal facilities. The TV adopts a totally different approach by maximizing the daily use of a single operant box by many animals. Because there is a single behavioral box per TV, the equipment costs as well as the space occupied are greatly reduced. Additionally, maintenance and troubleshooting become easier. Human errors caused by manual assignments are removed from the equation, and all parameters related to training protocols are logged and reproducible. Importantly, errors caused by the system (software or hardware) can be traced back easily as the TV automatically stores the system’s data, including video, together with the details of the training protocol and the identity of the subject.

Beyond multi-box systems, an increasingly adopted strategy is the use of home cage systems that enable 24/7 autonomous functioning without human intervention. While the majority of available home cage systems focus on monitoring ethological behaviors and social interactions^26,32,41,50,59,60^, some have also been designed to train and test rodents on trial-based tasks, either directly inside the home cage^23,25–28,30,37,61^ or in a separated operant box accessed via automatic doors^24,29,35,38,62^. Training directly inside the home cage limits modularity and adaptability of the system across behavioral paradigms. In addition, interference from cage mates during task execution may distract animals and impact performance. Among the systems that train freely-moving animals in a physically separated operant box, the only open-source, multi-animal platform to date is the Mouse Academy. This system is based on Python code running on a PC or Raspberry that, by a series of RFID antennas and two doors, controls access to a separate operant box^24^. Despite the similarities, the TV offers a number of advantages over the Mouse Academy: it has both mouse and rat versions (Supplementary Fig. S1); it has a double detection system in the corridor based on RFID and video analysis which minimizes the probability of two animals entering the operant box (Supplementary Fig. S2); it measures activity in the home cages (Fig. 4); it is equipped with a user-friendly GUI for remote access and monitoring of task progress within and across sessions (Supplementary Fig. S3); it has been extensively tested over multiple tasks (Fig. 3); it is equipped with a number of sensors for the close supervision of the animals’ welfare such as a scale for tracking weight changes, humidity and temperature probes and a system of alarms that allows a prompt response to any unexpected event. A detailed comparison of the different home cage training systems can be found in Supplementary Table 1.

The TV is specifically designed for complex cognitive tasks that typically require extensive manual training over weeks or months and considerable experimenter expertise. The system can achieve complete automation of the training process by an adaptive training algorithm that, based on the performance of each individual animal in each session, decides which task stage and parameters to use in the next one (Fig. 1c). Together, the real-time alarm notifications and automatic daily performance reports make the platform suitable for long-term autonomous operation while still allowing detailed remote supervision, making it ideal for longitudinal studies such as pharmacological testing or aging. Moreover, this full automation enables a clear division between routine maintenance tasks (cleaning, water refill, calibration, etc.), which require minimal expertise and detailed assessments of animals’ performance, which can be addressed by skilled researchers remotely. By reducing repetitive manual work, the system also alleviates the burden on trainees and supports healthier working conditions. Finally, automation improves reproducibility across groups and opens opportunities for inter-laboratory collaboration, for example, enabling researchers to train animals remotely in partner institutions.

Using the TV, we showed that animals were able to successfully engage in three cognitively demanding tasks of varying complexity and spanning distinct cognitive functions, including decision-making, short-term memory, attention, perceptual discrimination and reinforcement learning. The ability of animals to maintain high accuracy and complete large numbers of trials across these paradigms (Fig. 3j-k and Supplementary S6a) indicates that autonomous, self-paced training does not compromise task engagement or performance, even for tasks traditionally considered challenging to implement without close experimenter supervision. Notably, the number of trials per day performed by mice in the TV was comparable to animals trained manually, although distributed in multiple shorter sessions (Supplementary Fig. S7). Considering however that mice in the TV can work every day (including weekends), the final trial count after a few weeks of training rapidly exceeds those obtained with manual training. Importantly, task performance in self-trained animals was comparable to, if not better than the animals trained manually (Supplementary Fig. S7), consistent with previous studies using automated systems^25,27,31,37^. In addition, animals effectively regulated their own training schedules, engaging with the task most of the time spent inside the operant box (Fig. 3f), without experimenter-imposed timing or deprivation schedules (e.g., single daily sessions at fixed times in manual training). This self-initiated mode of training may better capture natural fluctuations in motivation and arousal, which influence cognitive performance but are often ignored in conventional training paradigms. Overall, the successful implementation of multiple tasks highlights the versatility and modularity of the TV.

Beyond task performance and engagement, the TV enables continuous collection of complementary behavioral measures, such as the inter-session intervals, circadian activity, putative sleep periods, and number of daily sessions, enabling detailed analyses on how different temporal patterns can shape task behavior. Such measurements provide access to behavioral variables that are largely inaccessible in standard laboratory paradigms, in which testing is typically restricted to short, fixed daily sessions. Moreover, by continuously tracking animals’ location in the home cages, the TV enables the assessment of individual and social preferences and their relationship with cognitive abilities. Although few studies have explicitly linked home cage behavior and individuality to cognitive functions^38^, growing evidence suggests that individual differences in activity and social interaction can emerge spontaneously and remain stable over time. Capturing these dimensions alongside cognitive performance provides critical context for interpreting variability in learning, motivation and strategy, and opens new avenues for studying how ethological traits interact with cognition under minimally constrained conditions.

We adapted the TV for rats, enabling cross-species comparisons. Several studies have previously addressed differences between mice and rats in task performance. In general, both species are capable of learning cognitively demanding tasks with high accuracy, although mice sometimes show slightly lower performance^63^. The major differences are found during task acquisition, with rats often learning faster^64–67^, and exhibiting more stable performance over time^63,66,67^. Additionally, important differences in the strategies used to solve the tasks have been described^63,65,68^. Our results show that both species used the setup similarly, although some differences in motivation were observed: rats performed fewer sessions per day than mice with similar group sizes (Fig. 2f), and spent less time engaged in the task (Fig. 3f). These differences may reflect greater persistence in mice. Opposite to our results, previous studies using touchscreen-based operant paradigms observe a larger number of daily trials in rats^66^ and higher impulsivity^69^. However, our observed differences could also result from factors such as reward size or sucrose concentration not being proportionally scaled between species and across studies. Further testing is required to clarify these differences.

The TV is designed to be extended and integrated with other systems. As we have demonstrated with the Eco-HAB^41^, other open-source systems can be integrated as extensions, such as modules for self head-fixation^16,25,61^, highly precise home cage monitoring systems^60^, or wireless battery-free optogenetics^31,70^. The latter has already been integrated with the TV and we are currently collecting preliminary data. Although some brain recording or manipulation techniques cannot be adapted to the TV (e.g., cables or batteries required), the system can still be used in hybrid mode. In such cases, animals undergo manual sessions on days when recordings or manipulations are performed, and automatically train on periods when they are not tested (e.g., weekends). The TV can be configured to block individualized access to the operant box during user-defined periods, ensuring high motivation for testing sessions. We have successfully used this hybrid approach across multiple experiments (e.g., electrophysiology, intracranial infusions, tethered optogenetics, and pharmacological manipulations). Additionally, with minor adjustments in the corridor access protocol (enabling the entrance of two animals simultaneously while tracking their identities) the TV is ideal to test social behavior using competitive^71^, pro-social^72,73^, and cooperative^74,75^ multiplayer tasks. Moreover, the system could be easily extended for supporting the access to more than one operant box running different paradigms, which is necessary for studying the task specificity of certain brain areas or manipulations^76^.

The TV is strongly aligned with improving the wellbeing of animals used in laboratory research. It improves animal welfare by minimizing their transportation and handling (i.e. *Refinement*). It allows animals to live in enriched environments in medium-sized groups (10-15) so that they can display rich social interactions. Importantly, animals are not technically deprived of food or water, as commonly done during behavioral experiments. Instead, they have continuous access to water with CA^49,57^ and can forage for more palatable water by doing the task at any time of the day. The TV incorporates individual tracking of animal variables including weight, water intake, locomotor activity and task performance, paired with an alarm system which enables experimenters to react promptly to any circumstance that may be causing animal suffering, (e.g. high temperatures, low subject weight or water intake; see Supplementary Table 2). Moreover, because the system can run non-stop over months (Fig. 2a-b, h), the amount of data obtained from each animal can be orders of magnitude larger than in manual behavioral experiments, which results in the reduction of animal group sizes. Overall, the operational principles and monitoring capabilities of the TV raise considerably the current standards of animal welfare.

Finally, the TV represents a powerful tool for translational research. Despite major advances characterizing neural circuits underlying higher-order cognitive functions, many psychiatric and neurodegenerative disorders still lack effective treatments, and the success rate of drug development has been disappointingly low over recent decades^77–80^. This is partially due to the complexity of the disorders, but also because of the poor translational validity of preclinical models: most of the compounds that show efficacy in rodents ultimately fail in clinical trials. A key limitation is the lack of robust behavioral biomarkers that could be measured longitudinally across species to inform diagnosis, prognosis, and treatment response. Fully automated systems that combine continuous testing in cognitive tasks adapted from humans with monitoring of spontaneous home cage behavior offer a promising approach to address this challenge^81^. By enabling long-term assessment of behavior under conditions that more closely resemble clinical study designs, these platforms allow the evaluation of chronic pharmacological interventions and disease progression over extended timescales with large and robust datasets. The TV integrates these capabilities offering a promising approach for improving the translational relevance of preclinical behavioral studies, bridging the gap between rodent models and human cognition.

In conclusion, the TV is an affordable, versatile, effective, convenient, controllable and respectful system for rodent training. It maximizes the use of the operant box, reduces human involvement, produces high-quality data across complex cognitive tasks, and enables longitudinal studies with continuous monitoring of cohort animal behavior under improved welfare conditions.

## Methods

### Subjects

A total of 115 mice were trained in groups using the Training Village, including 87 C57BL/6J, 4 Gad2-Cre, 7 Pvalb-Cre, 7 Pvalb-Cre Grin1 conditional knockout, and 10 TRAP2 x Ai14 mice. Animals were trained in groups on different cognitive tasks: 3-choice visuospatial delayed response task (92 mice, 8 groups), 2-choice perceptual discrimination (10 mice, 1 group), and 2-armed bandit task (13 mice, 2 groups). In addition, 6 female Long-Evans rats were trained in the visuospatial task using an adapted version of the TV. Some groups decreased the number of subjects over time. Detailed information about the groups is provided in Supplementary Table 3. Animals had *ad libitum* access to food and water with citric acid (2-4%) in their home cages (except Groups 2 and 3 which received water only during the task). Inside the operant box, they received 10% sucrose water as a reward. Animals were 6–12 weeks at the beginning of the experiment and were group-housed under controlled conditions (22±2°C, 12 h light/dark cycle, lights on at 8 a.m.). Body weight was monitored daily, and animals dropping below 80% of baseline were temporarily removed and given free water access until recovery. Environmental enrichment (igloos, running wheels, plastic tubes, chewing blocks, nesting material) was provided to reduce stress. All mouse procedures were approved by the Animal Experimentation Ethics Committee (CEEA) of the University of Barcelona under project 33/20. Rat procedures were approved by CEEA of the Universitat Autònoma de Barcelona (UAB) under project CEEAH 4907-CEEA-UAB.

### Microtransponder Implantation

To individually identify animals in the TV, mice were subcutaneously implanted with glass-covered microtransponders (12 mm length, 2.12 mm diameter, RFIP Ltd) under brief isoflurane anesthesia before the training started. Microtransponders emit a unique animal identification code when within the range of RFID sensors.

### Hardware Architecture

The Training Village consists of a sorting corridor connecting the animals’ home cages to a custom operant box. The current system implementation uses a Raspberry Pi 5 as the central controller for all sensors, actuators, and cameras, running custom open-source Python software. Earlier cohorts in this study were tested using a first-generation implementation based on a desktop PC with USB cameras, an Arduino controlling the corridor, and fewer integrated features. The transition to the Raspberry Pi architecture provided substantial improvement in stability and reliability, and greatly simplified system installation. These improvements arise from the device’s suitability for continuous 24/7 operation, native camera integration, low power consumption, low cost (which facilitates the replacement), and direct compatibility with Python-based control software. The system includes the following components controlled by the Raspberry Pi: an RFID reader for animal identification (ID Innovations ID-12LA or Eco-HAB); a load cell with an ADS1114 ADC for body weight measurement; two MG995 servo motors to control corridor doors; an SHT31 temperature–humidity sensor for environmental monitoring; two overhead IR-sensitive cameras (Raspberry Pi Camera Module 3, 12 MP wide NoIR); and a programmable illumination system with white LEDs for day–night cycle simulation and continuous infrared lighting for video acquisition.

All peripheral connections, except the cameras (which use dedicated CSI connectors), are routed through a custom Hardware Attached on Top (HAT) board that interfaces with the Raspberry Pi GPIO pins. The HAT provides integrated connectors for all peripherals and accepts a 5 V external power supply to drive the RFID reader and servo motors. The board incorporates an LDO (voltage regulator) essential for reliable RFID operation, as well as a signal-conditioning circuit consisting of a low-pass filter, an amplifier, and an analog-to-digital converter (ADS1114) for the load cell. In addition, it provides connectors for an auxiliary servo motor and programmable LEDs. The system can alternatively be operated without the HAT by connecting components directly to the GPIO pins. However, this configuration requires a linear 5 V power supply (or external regulator) and an external HX711 amplifier/ADC module for the load cell, and results in less robust connections.

### Corridor Construction

The corridor was fabricated from PLA 3D-printed components and laser-cut white acrylic panels. Its geometry was iteratively optimized to ensure reliable single-animal passage while minimizing the physical footprint. The default white coloration maximized visual contrast for dark-coated rodents; this scheme can be inverted for light-colored animals. Detailed dimensions are provided in Supplementary Fig. S1a.

### Operant Box

Cognitive testing was performed in a custom-made operant chamber made of white acrylic. The operant chamber was inside a light and sound-attenuating box (medium-density fiberboard with acoustic foam), equipped with a fan, overhead lighting, and an infrared overhead camera for task performance monitoring. The chamber design was adapted to the specific task and species. For the visuospatial delayed-response task in mice, the arena consisted of an elevated maze with two opposed triangular areas connected by a rectangular corridor (25.5 x 4 cm) (Fig. 3a). The entire arena was elevated 20 cm to avoid walls at the edges of the corridor and thus ensure clear visibility of the touchscreen, and prevent mice from exiting the arena. The larger triangular area contained an infrared touchscreen (43 x 25 cm, ePanel Closed 19”, EOS Iberica) used to display stimuli and record mouse poke responses. A black mask with three 6x6 cm response windows (13 cm spacing) covered the screen. The reward port (Sanworks) was located in the opposite triangular area (57 cm total maze length). The connecting corridor and the reward port were equipped with infrared beams to detect animal crossings. A piezoelectric buzzer above the touchscreen provided auditory feedback for correct or error responses. To prevent mice from escaping the chamber in case they jumped from the platform, two white acrylic side walls (60 x 50 cm) were installed from the sides of the touchscreen to the smaller triangular area. For rats, dimensions were scaled up (Fig. S1a). For the perceptual discrimination and two-armed bandit task, a rectangular arena (32 × 35 cm) was used, containing three custom-made reward ports with photo beams located on the larger wall opposite to the corridor (2 cm above the floor, with the center of each port 4 cm apart). Ports were illuminated, and rewards were delivered only through the side ports. The operant box was mainly controlled by Bpod (Sanworks), except for the touchscreen, which was controlled by the Raspberry Pi.

### Home Cages

Standard housing cages for mice or rats were manually perforated and connected with transparent acrylic tubes (inner diameter 3 cm, outer 3.4 cm for mice; inner 8.4 cm, outer 9 cm for rats) and covered with stainless-steel bar lids. Typical cage dimensions for mice were 22 x 22 x 15 cm or 22 x 47 x 15 cm for XL cages (Panlab). For rats, three different cages were used simultaneously in the same setup (46 x 40 x 40 cm, 48 x 38 x 21 cm from Tecniplast, and 71 x 46 x 31.5 cm from Ferplast). In groups 4-7, home cage activity was monitored using six circular RFID antennas surrounding the connecting tubes (∼5 cm from the edges). All the antennas were connected to a dedicated reader unit, which transmitted the data to the Raspberry Pi (reader and antennas provided by the Laboratory of Emotions Neurobiology, BRAINCITY, Warsaw).

### Software Architecture

The Training Village software is written in Python, runs continuously on a Raspberry Pi 5 under Raspberry Pi OS (Debian-based) and is fully open-source (https://github.com/BrainCircuitsBehaviorLab/village). The software monitors animals in their home cages, regulates access to the operant chamber, launches behavioral tasks, logs data, and manages remote communication. The system operates through a state machine in which each state defines the current system condition (e.g., waiting, manual mode, running a task, synchronizing data) and the permissible transitions (Fig. 1c, Supplementary Fig. S2). A detailed description of all states is provided in the online documentation.

### Graphical User Interface

The software includes a GUI that provides user-friendly interaction and real-time monitoring of the TV system. The GUI is organized into seven screens accessible via menu buttons: MAIN, MONITOR, TASKS, DATA, WATER_CALIBRATION, SOUND_CALIBRATION, and SETTINGS.

MAIN is the default screen in which the Raspberry Pi performs no active rendering. MONITOR displays the system status, including live video streams from both the corridor and operant box cameras; central control buttons provide quick access to key functions such as door control, scale readout, and temperature monitoring, while recent system events are logged in the lower panel (Fig. 1c). TASKS lists all available behavioral protocols and enables manual execution of them (Supplementary Fig. S3d). DATA provides an interactive interface for exploring saved experimental data, along with a list of the training parameters for each subject that are automatically updated based on user-defined criteria and animals’ performance (Supplementary Fig. S3c). Users can easily edit these parameters using this tab. The DATA tab also includes user-configurable plots for visualizing task performance (Supplementary Fig. S3e). WATER_CALIBRATION and SOUND_CALIBRATION provide interfaces for calibrating water delivery and audio output, respectively. SETTINGS displays an interactive list of all configurable parameters.

### Animal Detection Algorithm

A lightweight, CPU-efficient algorithm processes the corridor camera feed in real time. Each frame is binarized using a fixed luminance threshold, with pixels classified as black (below threshold) or white (above threshold). Black-pixel counts within predefined regions of interest determine whether each zone is empty, occupied by a single animal, or occupied by multiple animals (Supplementary Fig. S2b–c). This approach is computationally robust, high-speed, and suitable for continuous 24/7 sorting operations. The classification polarity can be inverted (bright animals against a dark background) when needed. Although the dataset presented here does not include software capable of tracking animals’ precise position in real time—either in the corridor or in the operant box—we are currently testing such functionality in new cohorts and will be incorporated in new versions of the TV. This novel tracking algorithm estimates the animal’s two-dimensional position (x, y) on each video frame. It is based on a lightweight OpenCV pipeline using contour detection (findContours). The processing time per frame is approximately 7–8 ms, which allows real-time tracking while maintaining a stable acquisition rate of 30 frames per second, with only about 1% of frames exceeding the nominal 33 ms frame duration. Tracking is performed at the camera’s default resolution (640 × 480 pixels). Although higher resolutions or frame rates are possible, these configurations increase the proportion of frames for which processing time exceeds the desired frame interval.

### Remote Access and Monitoring

Despite being a highly autonomous system, the TV requires supervision to ensure optimal performance, especially at the beginning of an experiment when animals are still not familiarized with the system or the task. Close monitoring and adjustment of some parameters during the first week after setting up a new system are always required (e.g., camera luminance thresholds for accurate animal detection). Once stabilized, the system remains remarkably steady over long periods. Nevertheless, unexpected issues that require fast attention may arise. To facilitate this, the TV is designed for remote operation and continuous monitoring.

Users access the system via standard remote desktop or VNC connections. An integrated Telegram alarm system delivers real-time notifications and enables remote status queries through a dedicated chat group managed by a Telegram bot (see Supplementary Table 2 for more details about the notifications). Twice daily, at each light-cycle transition, the system automatically evaluates the status of all animals and generates a report containing detection counts, session counts, water intake, and mean body weight for each subject. An alarm is triggered if any subject falls below predefined thresholds for any of these variables. In addition to these alarms, the software continuously monitors for errors and unexpected conditions, including sensor malfunctions, corrupted files, pixel detections in forbidden regions, task execution failures, settings update errors, and sessions with no recorded trials. Every detected error triggers an immediate Telegram alarm. The system is designed to handle all anticipated failure modes, resulting in an extensive library of alarms; while many may never be triggered, others may occur once every several months. A summary of the system alarms can be found in Table 2. A complete list of alarms and recommended corrective actions is available in the online documentation. To further ensure system uptime, an external monitoring service (healthchecks.io) receives an hourly heartbeat signal from the Raspberry Pi and automatically issues an alarm if the signal is missed.

While most problems can be resolved by connecting remotely to the TV (e.g. two subjects entered simultaneously into the operant box) others may require on-site intervention (e.g. loss of network connectivity). Importantly, the availability of CA water in the home cages eliminates the need for immediate intervention if an animal fails to consume the minimum required amount of water from the operant box. Because animals can safely rely on CA water for 2–3 days, any issue can typically be addressed on the next regular workday. If preferred, traditional water restriction is fully compatible with the TV, but it requires closer daily monitoring and rapid responses to any system malfunction.

### Cognitive tasks

Three cognitive tasks were tested in the TV. Given its modular architecture, tasks shared the same software, corridor, and home cages, and only the operant box, task code, and settings file were changed. ***Visuospatial delayed-response task:*** A three-alternative forced choice (3AFC) paradigm in which animals had to memorize a stimulus position and navigate to the corresponding location. The task was structured as follows: A visual stimulus (white vertical rectangle 30 x 60 cm on a black background) was displayed on a touchscreen at one of three possible positions (Left, Center, and Right), randomly selected from a uniform distribution. As the animals ran through a corridor toward the screen, the stimulus disappeared at different points along the corridor, creating a variable mnemonic delay and four trial difficulty conditions: *Easy*, the stimulus remained visible until the mouse touched the screen (0 s delay); *Medium*: The stimulus was displayed until the mouse reached the end of the corridor (∼2 s delay for mice, ∼0.8 s for rats); *Hard*: The stimulus offset occurred when the mouse crossed the late middle of the corridor (∼2.3 s delay for mice, ∼1.5 s for rats); *Extreme*: The stimulus was only visible at the very beginning of the corridor (∼3.3 s delay for mice, ∼2.6 s, for rats). Subjects reported the location of the stimulus by nose-poking the screen at the remembered position. Correct responses were signaled by a 15 kHz auditory cue, informing animals to run back opposite to the screen to collect a 10 μl reward of 10% sucrose water. Incorrect responses triggered a 4kHz auditory signal and the illumination of the operant chamber, ending the trial without a reward. Omitted responses were also cued with the chamber lights onset and not rewarded (Fig. 3a). After errors or omissions, animals had to return to the water port area to initiate a new trial. ***Evidence discrimination task, visual version:*** A 2AFC paradigm in which mice discriminated the relative intensity between two light stimuli. Trials were self-initiated by a nose poke at the illuminated center port. Upon initiation, the center LED turned off, and both side ports were illuminated simultaneously with different intensities. Mice were required to maintain fixation in the center port for at least 500 ms before choosing the side port with the brighter stimulus. Fixation breaks restarted the trial. Across trials, three difficulty levels were randomly interleaved according to the luminance ratio between ports: *Easy* (5), *Medium* (2.5), *Hard* (1.25). Only correct responses were rewarded with 2 μl of sweetened water (Fig. 3b-c). Each trial was followed by a 1s inter-trial interval (ITI), after which the center LED was illuminated again to signal the next trial. ***Auditory version:*** The task structure was the same but the stimulus was a sound that consisted of a stream of 30ms pure tones presented at 100Hz. One of two octaves (5 to 10kHz, or 20 to 40kHz) was selected as the target octave to indicate the correct side port. Individual tones were randomly drawn from 6 possible frequencies on each octave. The difficulty of sound discrimination was controlled by varying the proportion of tones from each octave, defined as the stimulus evidence: *Easy* (0.98 vs 0.02), *Medium* (0.82 vs 0.18), *Hard* (0.66 vs 0.33). Sound amplitude was randomly selected between 60 and 80 dB. ***Probabilistic two-armed bandit task:*** Animals had to track which of the two sides was more likely to deliver a reward. Trials were initiated when mice poked the illuminated center port. Immediately afterward, both side ports were illuminated, indicating which side to choose. The reward probability for each side depended on the current block, requiring mice to infer the higher-probability side based on recent reward history (Fig. 3c). Animals received 3 μl of sweetened water as a reward. Four difficulty levels were used, defined by the reward probability of the high-probability side: *Easy* (0.9), *Medium* (0.8), *Hard* (0.7), and *Extreme* (0.6-0.5), with the opposite side having the complementary probability. Reward contingencies were reversed periodically across blocks, which lasted between 25 and 55 trials randomly. After a choice, animals experienced a variable ITI drawn from an exponential distribution (mean = 5 s, range = 0–30 s).

### Data processing and analysis

Different datasets were preprocessed for analysis. To evaluate the general system usage (Fig. 2, Supplementary Fig. S4-5), a DataFrame containing all corridor events was used. The DataFrame had one row per event, and each row included the event timestamp, subject identity, group number, whether an entrance attempt was successful, and—if successful—the corresponding session start and end times. Failed entrance attempts were labeled with the specific failure type (multi-animal or refractory). It is important to consider that some of the multi-animal attempts are also occurring in the refractory interval, but their exact proportion cannot be determined because once a multi-animal attempt is detected, the TV software no longer checks whether the refractory interval has elapsed. The occupancy of the operant box was calculated as the percentage of time in a day (24h) in which the box was occupied by one mouse.

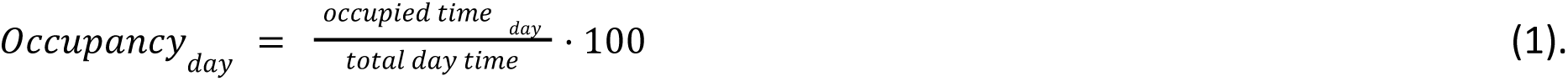

The Gini coefficient (*G*) was used to quantify inequality in box usage across subjects of the same group. Animals were sorted by occupancy, and the cumulative fraction of subjects as a function of the cumulative fraction of occupancy L(*p*) was calculated. *G* was then computed as:

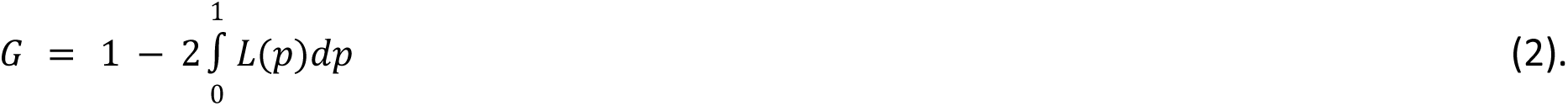

A Gini coefficient of 0 indicated perfectly uniform occupancy across subjects, whereas 1 indicated that a single subject accounted for all occupancy. Using this dataset, we also quantified the number of failed entrance attempts per successful session and the number of sessions performed by each subject within a day.

For the cognitive assessment (Fig. 3), we used a DataFrame containing one row per trial. Columns included session date and number, subject number, group number, trial duration, trial outcome, trial difficulty, and task type. To determine the engagement of each trial, for each subject, we computed its mean trial duration and classified trials as highly engaged (<0.5σ below mean), medium engaged (−0.5σ to +2σ), or disengaged (>2σ). For most analyses, highly and medium engaged were clustered together. Because the 2AB had a variable forced inter-trial interval, this epoch was removed from the trial duration for engagement analysis. Accuracy was defined as the fraction of correct trials over the total number of valid trials (trials in which the animal responded within the response window). For the 2AB task, a trial was classified as correct when the animal poked the side associated with the higher reward probability, even if no reward was delivered on that particular trial. For analyses assessing engagement (Fig. 3h) or trial duration (Supplementary Fig. S6d) as a function of trial index, we controlled for differences in total trial number across sessions. Without this control, higher trial indices would be represented only by the longest sessions, which tend to have higher overall engagement. To avoid this confound, we restricted the analysis to sessions with a total trial count at or above the median across all sessions, and computed the metrics only up to the median trial number. This ensured that every data point in the plot was supported by the same set of sessions. Additionally, for the 3AFC task in mice we excluded the first 12 trials from this analysis, as they constitute an initial warm-up period consisting only of easy trials designed to promote early task engagement. This warm-up was not present in the other tasks. Lastly, for the comparison between automating and manual training in task performance (Supplementary Fig. S7) we only included sessions after animals completed the training protocol. Notably, this comparison was not conducted between two groups of equal size trained in parallel under identical conditions; therefore, results should be interpreted as reflecting general trends rather than definitive evidence of equivalence.

For the home cage activity assessment (Fig. 4), RFID antenna lectures were processed in a DataFrame with one row per detection, with the timestamp, antenna number, and subject identity. Home cage occupancy was inferred from the sequence of antenna detections: when two consecutive antennas corresponding to a given cage were activated by the same subject, the subject was considered to be inside that cage. Cage changes were also extracted and used for Fig. 4c-d. These were defined as successive detections of antenna pairs within the same corridor connecting two cages by the same subject occurring within a short time interval (< 1 minute). Finally, to ensure a fair comparison between the task and no-task conditions in Fig. 4d, periods during which each animal was inside the operant box were excluded from the no-task condition, as animals cannot perform cage changes while engaged in the task. Some of the variables described previously were separated by hour or by day/night condition.

## Author Contributions

Conceptualization: J.dR., R.M., B.S.; Software: R.M, H.M.V., B.S.; Hardware: R.M. (mice), D.P. and H.V. (rats); Experiments: B.S. (3AFC task mice), H.M.V. (2AFC task), C.B. (2AB), H.V, D.K. and A.T. (rats); Data Analysis and interpretation: J.dR., B.S., R.M., J.R., H.M.V.; Documentation and website: R.M., J.R., H.M.V., B.S.; Writing: B.S., J.dR.; Reviewing and editing: H.M.V., R.M., A.T. These authors contributed equally: B.S. and R.M.

## Data and code availability

Training Village software, hardware designs, assembly instructions, and user documentation are available in a GitHub repository (https://github.com/BrainCircuitsBehaviorLab/village). Data supporting the figures will be made available upon peer-reviewed publication.

## Acknowledgments

The authors would like to thank Emma Condon, Andrés Crespo, Eva Carrillo, and Juliette Merrheim for their support during the experiments, Eric D’Andria for technical help during the development of the Raspberry Pi HAT, Javier Garrido for his support with the 3D designs, Juan Garrido for assistance in developing the pilot versions of the TV, Ewelina Knapska and Aljcia Puścian for providing an Eco-HAB system.

Funding was provided by the Spanish State Research Agency (AEI) (grants PID2021-126698OB-I00 with EU NextGeneration funds; PCI22_135057-2 with EU NextGeneration funds; PID2023-150497OA-I00), the European Research Council (grant ERC-2022-POC2_cutoff-101101098-MOUSEVILLAGE and ERC-CoG-101043986-UNIPROB), the National Institutes of Health (award number 1R01MH132172-01), and the Cellex Foundation. Part of this work was done in the Centre Esther Koplowitz.

## Abbreviations

2AFC: Two-alternative forced-choice
2AB: Two-armed bandit
CA: Citric acid
GUI: Graphical user interface
ISI: Inter-session interval
RFID: Radio frequency identification
TV: Training Village.

