## Supplementary figures and tables for "The Training Village: an open platform for continuous testing of rodents in cognitive tasks"

### Extended data

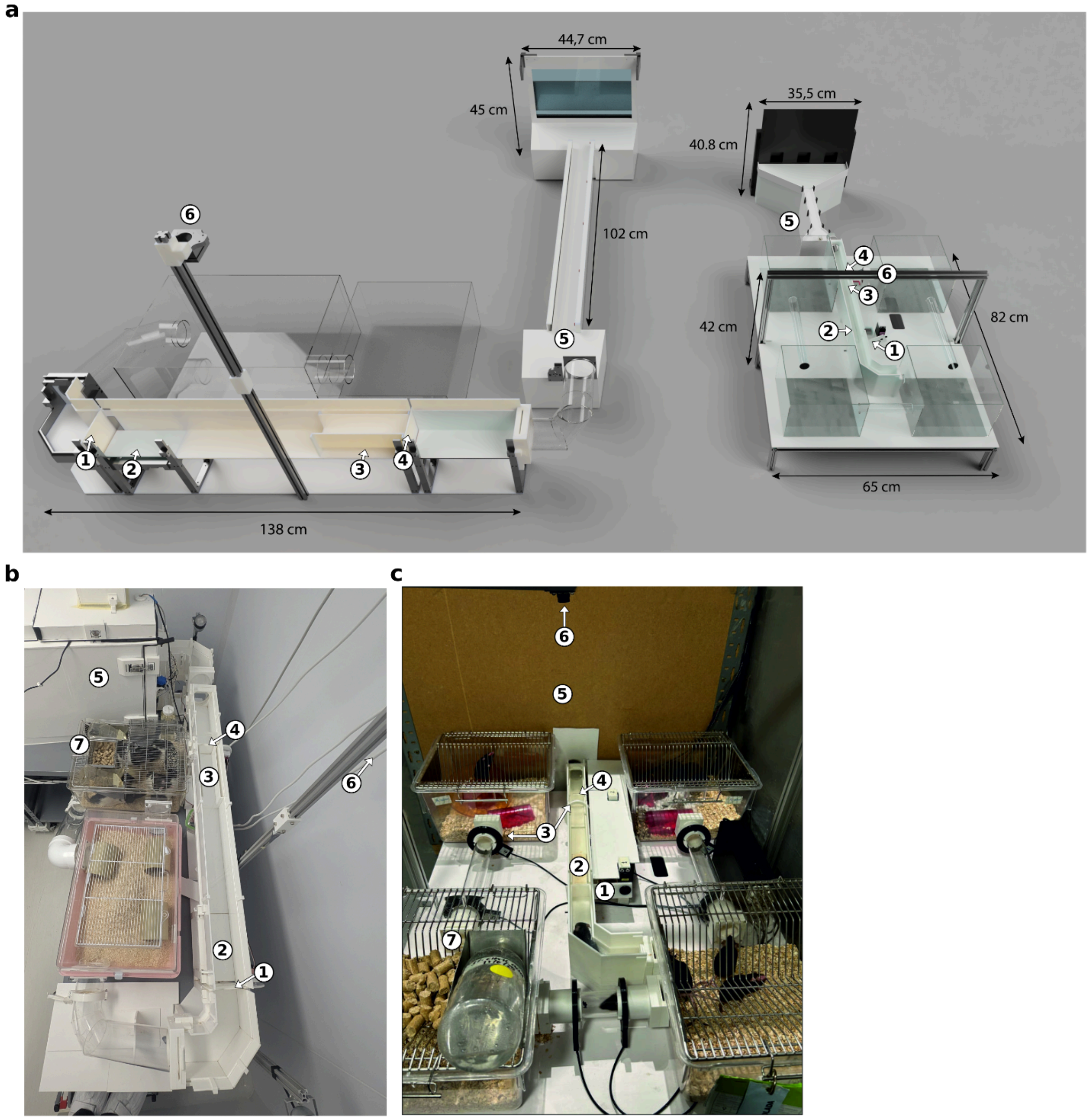


**Figure S1: Training Village design. a,** 3D renderings of the rat (left) and mouse (right) Training Village including the home cages (rendered in glass and empty for easy visualization), the corridor, and the interior of the operant box featuring the touchscreen maze used for the 3AFC task (Fig. 3a). The isolation enclosure of the operant box and walls of the rat's corridor have been removed to allow visualization of the setup. Numbers indicate the different parts: 1 door one, 2 scale, 3 RFID readers, 4 door two, 5 operant box, 6 IR camera, 7 food and citric water. **b-c,** Pictures of the rat TV (b) and the mouse (c) TV.

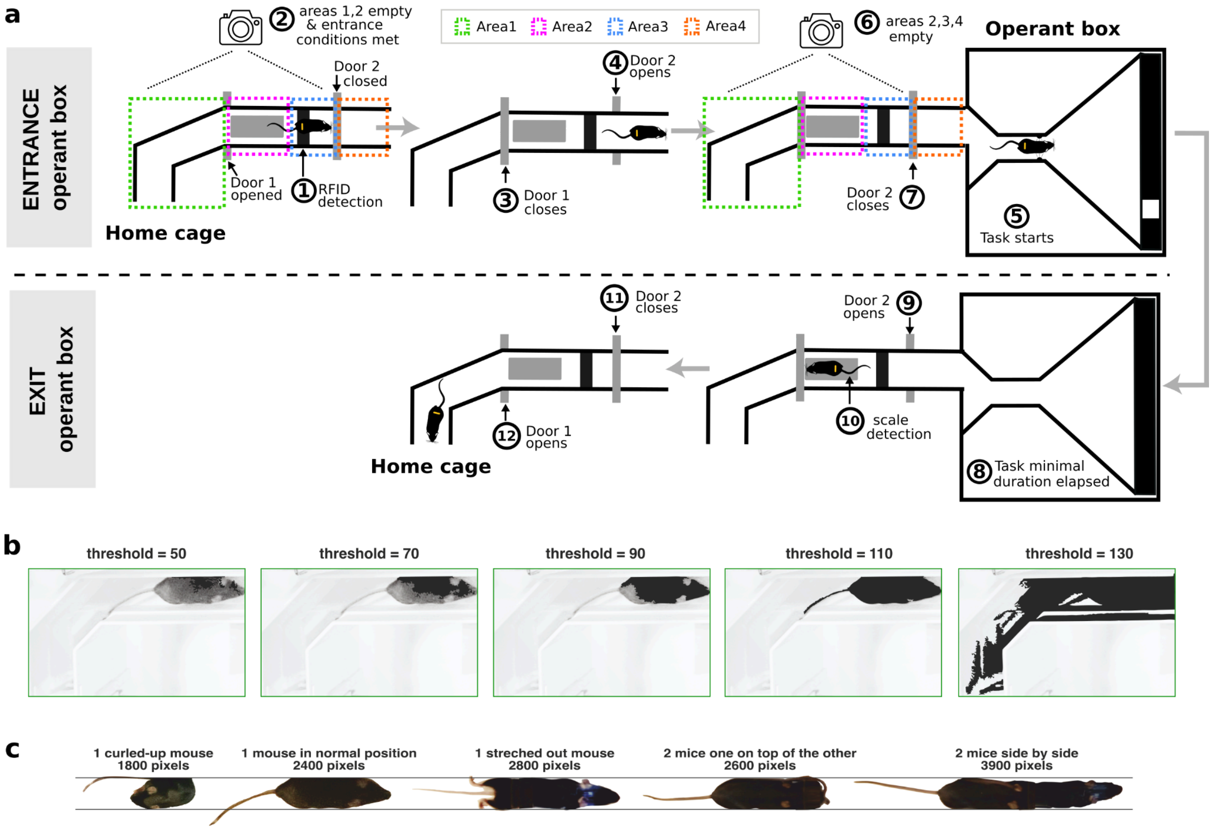


**Figure S2: Corridor system for granting individualized access to the operant box. a,** Diagram of the automated process to enter and exit the operant box in an individualized manner. RFID-tagged animals grouped in common home cages access the behavioral box through a corridor with two doors controlled by a double detection system composed of an RFID reader and video analysis. When the operant box is empty, Door 1 (near the home cage) is open and Door 2 (next to the RFID) is closed, allowing animals to enter the corridor. When a subject reaches Door 2, the system checks if there are more animals in the corridor by online video analysis of the pixels in the camera Areas 1 and 2 (green and pink boxes respectively). It also checks that Area 4 is empty (orange), a condition that must always be satisfied when Door 1 is open. Finally, the system verifies that there is only one animal in Area 3 (blue). If the animal is alone in the corridor, currently active (animals can be configured to be active or inactive on different days), and the refractory interval has elapsed, the system grants access by closing Door 1 and opening Door 2, after which the task starts. Once the animal is inside the operant box, Areas 2, 3, and 4 are detected as empty by the camera and Door 2 closes. This procedure guarantees that only the RFID-detected animal enters the operant box, and doors are safely closed without any mice being trapped. At the end of the minimal session duration, Door 2 is opened and the mouse can leave the box. When it exits, it is detected by the scale which stores its weight and triggers the closing of Door 2, followed by the opening of Door 1, allowing the mouse to return to the home cage. **b,** Video analysis algorithm used to detect animals in the corridor by binarizing frames (black and white) according to a fixed brightness threshold. Optimal calibration of the brightness threshold requires maximizing the number of detected pixels corresponding to the animal while minimizing false above-threshold pixel detections corresponding to parts of the corridor or elements outside (fourth example). **c,** Pixels detected by the algorithm in different frames with one or two subjects in the corridor and with different body positions. In these examples with a single mouse in the corridor (left three frames), the pixel area yielded values between 1800 and 2800 pixels. When two animals were present simultaneously, their combined area could be as small as 2600 pixels, producing overlap between the two distributions. To minimize cases in which two or more animals are classified as a single one and allowed to enter together (Type II errors), the threshold is set conservatively near 2500 pixels, even at the cost of increasing occasional false positives (Type I errors), where a single animal is temporarily classified as multiple and access is denied.

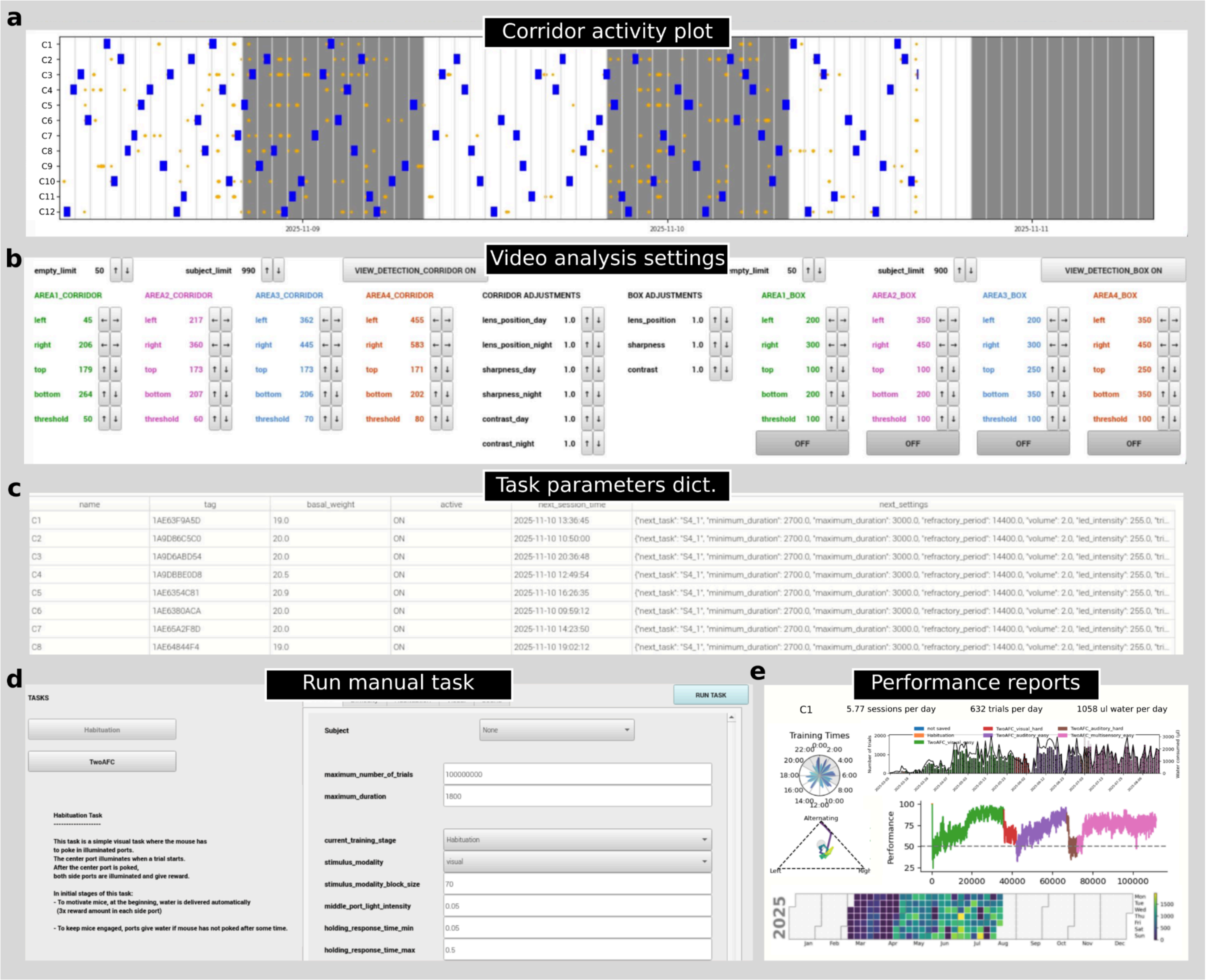


**Figure S3: Graphical User Interface of the Training Village.** The GUI offers multiple tabs to visualize animal activity, configure the system settings, manually run the behavioral tasks, and monitor subjects' performance, among other functions. Some tabs are shown here. **a,** Raster plot showing the activity of a group of mice (as in Fig. 2a). **b,** User-friendly, interactive panel to configure video analysis settings. **c,** Dictionary task parameters for each subject, easily editable by the user. **d,** Interface for manually running behavioral tasks, with quick access to key variables. **e,** Example performance reports for a mouse trained in the 2AFC perceptual discrimination task. These plots are configurable and should be adapted to the specific behavioral paradigm. *Top left*: Polar histogram of training times during the day. *Top right:* Number of daily trials across days, with colors indicating different task variations. *Center left*: Subject's systematic bias over time (e.g., repeating left, right, or alternating responses). *Center right*: Performance across trials, with colors indicating different task variations. *Bottom*: Calendar of training sessions with colors indicating the number of daily trials obtained.

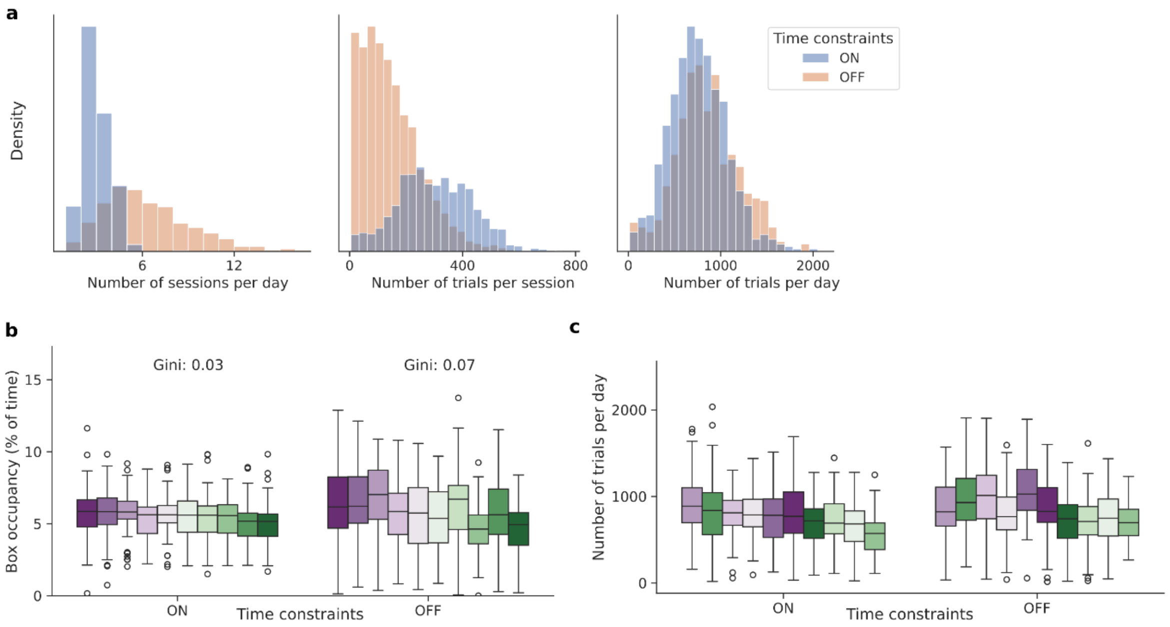


**Figure S4: Effects of temporal constraints in the usage of the operant box. a,** Histograms of the number of daily sessions per animal (left), number of trials per session (center), and number of daily trials accumulated per animal (right), computed separately for the two conditions (Time constraints ON vs OFF). Animals (Group 12, n = 10) were trained with temporal constraints, i.e., minimum session duration of 30 min and refractory interval of 4 h (Time constraints ON), and then tested in a condition without any time constraints (Time constraints OFF). In the OFF condition, animals could exit the box at any time after starting a session and could then re-enter at any time after exiting (provided there were no more mice in the corridor). **b,** Operant box individual occupancy for the two conditions (Time constraints ON vs OFF). Each box represents one subject and shows the median and IQR across days. Subjects are sorted by their occupancy in the ON condition. Although the distribution of time in the box is less uniform in the OFF condition (Gini ON vs OFF = 0.03 vs 0.07), it is still close to uniform. **c,** Number of daily trials per animal in the two conditions (Time constraints ON vs OFF). Same as in panel a. **c,** Analyses are based on 3956 “OFF” and 1906 “ON” sessions from Group 12.

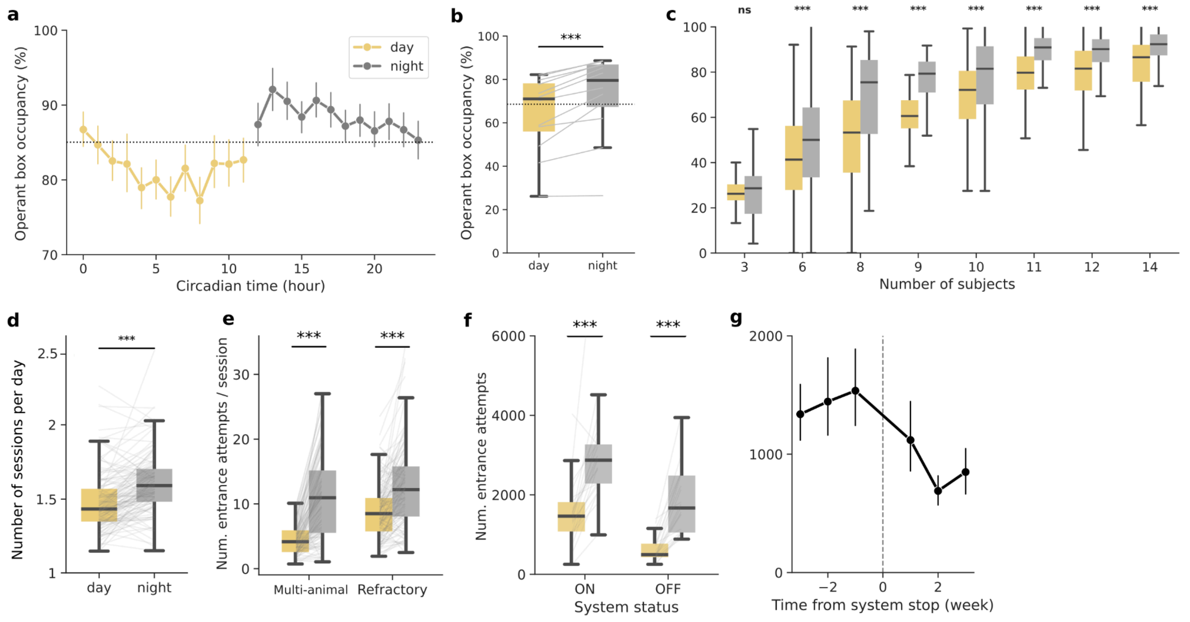


**Figure S5: Circadian effects on operant box usage. a,** Operant box occupancy over the 24-h day-night cycle. Dots represent the mean box occupancy per hour across sessions and subjects (circadian time). Error bars are 95% CI of the mean. The dashed line indicates the overall mean. **b,** Average box occupancy during day vs. night. Dots represent individual groups. The dashed line indicates the overall mean. Paired t-test $t$ = -5.8, $p$ < 0.001. **c,** Day and night box occupancy as a function of the group size. Asterisks indicate significance levels from paired t-tests comparing day vs. night (size 3: $t$ = -0.5, $p$ = 0.6; size 6: $t$ = -4.5, $p$ < 0.001; size 8: $t$ = -6.6, $p$ < 0.001; size 9: $t$ = -3.8, $p$ < 0.001; size 10: $t$ = -7.1, $p$ < 0.001; size 11: $t$ = -10.6, $p$ < 0.001; size 12: $t$ = -10.7, $p$ < 0.001; size 14: $t$ = -4.6, $p$ < 0.001). **d,** Average number of sessions in each light cycle. Paired t-test: $t$ =-7.3 , $p$ < 0.001. **e,** Number of failed entrance attempts depending on the light cycle. RM-ANOVA: attempt type $F(1, 120)$ = 30.9, $p$ < 0.0001; cycle $F(1, 120)$ = 278, $p$ < 0.0001; interaction $F(1, 120)$ = 17.6, $p$ < 0.0001. **f,** Number of failed entrance attempts depending on system status (on: normal access to the operant box; off: no access to the operant box and plain water *ad libitum* in the home cages). Repeated measures ANOVA: system status $F(1, 20)$=25.6, $p$ < 0.0001; light cycle $F(1, 20)$= 77.9, $p$ < 0.0001; interaction $F(1, 20)$=1.1, $p$ = 0.31. **g,** Number of failed entrance attempts aligned to the moment in which access to the operant box was discontinued (vertical line). Dots indicate subject mean counts per weekly bin, and error bars show the 95% CI of the mean. Boxplots show the median and IQR across groups (b), days (c), or subjects (d-f).

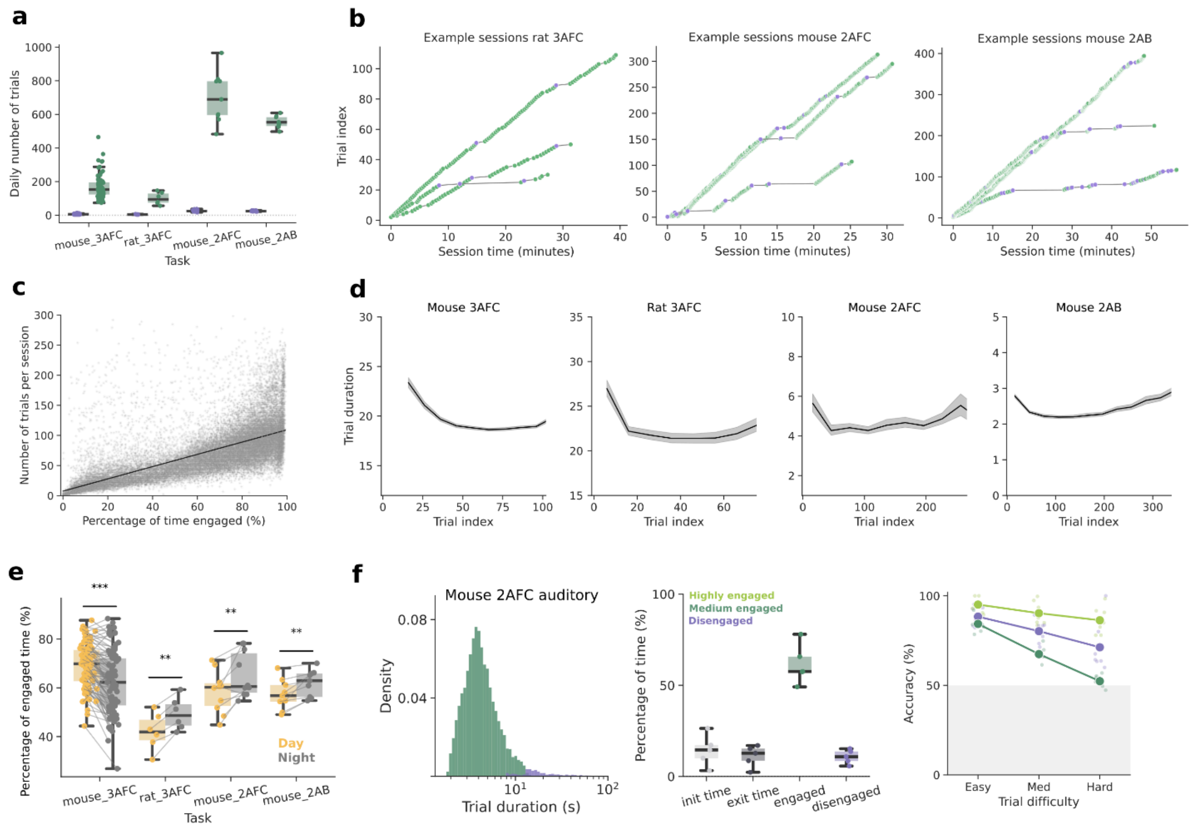


**Figure S6: Extended results of task engagement in the Training Village**. **a,** Averaged daily number of engaged vs disengaged trials performed by subject in the three tasks. Dots represent subjects, and boxes show the IQR with the median. **b,** Examples of animals' engagement along three example sessions in each of the three different tasks. As in Fig. 3e, green dots represent engaged trials and purple dots represent unengaged trials. **c,** Correlation between the number of trials per session and the percentage of engaged time within the same session. Each dot is a session. The line shows a linear regression fit (slope = 1, $p$ <0.001). Only mice from the visuospatial task were used. **d,** Trial duration as a function of trial index. Lines represent averages across bins (10-trial bins in 3AFC and 30-trial bins in 2AFC and 2AB). **e,** Circadian impact on the percentage of engaged time inside the operant box for the different behavioral tasks. Paired t-tests: Mouse 3AFC $t$ = 8, $p$ < 0.001; Rat 3AFC $t$ = -5, $p$ = 0.003; Mouse 2AFC $t$ = -3, $p$ = 0.01; Mouse 2AB $t$ = -4, $p$ = 0.002. **f,** Same plots as in Figure 3d, f, j, but computed for the auditory version of the 2AFC perceptual discrimination task.

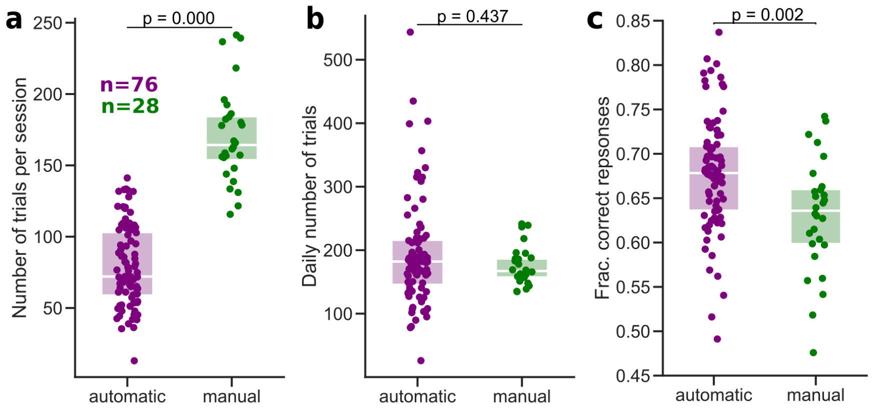


**Figure S7: Automatic training produces results comparable to manual training. a,** Average number of trials performed per session during automatic and manual training. **b,** Averaged daily number of trials performed under each condition. **c,** Task accuracy by training type. Dots indicate individual subjects. Boxplots show the median across subjects and the IQR. *P* values correspond to Mann–Whitney tests. Automatic training resulted in greater variability in daily trial counts and accuracy across subjects, likely reflecting individual preferences due to the increased flexibility of self-training.

| System | Specie | Automatic training algorithm | Group Housed | Home cage monitor | Isolated testing | Testing outside home cage | Touch screen | Alarms system | Open source |
|---|---|---|---|---|---|---|---|---|---|
| **Training Village** | Mouse/Rat | ✔ | ✔ | ✔ | ✔ | ✔ | ✔ | ✔ | ✔ |
| **Intellicage** (Kiryk et al., 2020) | Mouse | ✔ | ✔ | ✖ | ✖ | ✖ | ✖ | ✖ | ✖ |
| **Mouse Academy** (Qiao et al., 2018) | Mouse | ✔ | ✔ | ✖ | ✔ | ✔ | ✖ | ✖ | ✔ |
| **Phenosys**( Rivalan et al., 2017) | Rat | 🔶 | ✔ | ✖ | ✔ | ✔ | ✔ | ✖ | ✖ |
| **Home cage odor system** (Caglayan et al., 2021) | Mouse | 🔶 | ✔ | ✖ | ✔ | ✔ | ✖ | ✖ | ✖ |
| **Educage** (Maor et al., 2018) | Mouse | 🔶 | ✔ | ✖ | ✖ | ✔ | ✖ | ✖ | ✔ |
| **Home cage head-fixed** (Murphy et al., 2020) | Mouse | 🔶 | ✔ | ✖ | ✖ | ✖ | ✖ | ✖ | ✔ |
| **Automated 5CSRTT** (Birtalan et al., 2020a) | Mouse | ✔ | ✖ | ✖ | ✔ | ✔ | ✖ | ✖ | ✔ |
| **Automatic joystick task** (Bollu et al., 2019) | Mouse | 🔶 | ✖ | ✖ | ✔ | ✖ | ✖ | ✖ | ✔ |
| **ARTS** (Poddar et al., 2013) | Rat | 🔶 | ✖ | ✖ | ✔ | ✖ | ✖ | ✖ | ✔ |
| **Autocage** (Hao et al., 2021) | Mouse | ✔ | ✖ | ✖ | ✔ | ✖ | ✖ | ✖ | ✔ |
| **HABITS** (Yu et al., 2025) | Mouse | ✔ | ✖ | ✖ | ✔ | ✖ | ✖ | ✖ | ✔ |
| **Multiwhisker detection** (Bernhard et al., 2020) | Mouse | 🔶 | ✖ | ✖ | ✔ | ✖ | ✖ | ✖ | ✖ |
| **AutonoMouse** (Erskine et al., 2019) | Mouse | ✖ | ✔ | ✖ | ✔ | ✖ | ✖ | ✖ | ✔ |
| **Psibox** (Francis & Kanold, 2017) | Mouse | ✖ | ✔ | ✖ | ✖ | ✖ | ✖ | ✖ | ✔ |
| **Smart-Kage**(Ho et al., 2023) | Mouse | ✖ | ✖ | ✖ | ✔ | ✖ | ✔ | ✖ | ✖ |
| **Souris City** (Torquet et al., 2018) | Mouse | ✖ | ✔ | ✔ | ✔ | ✔ | ✖ | ✖ | ✖ |
| **Aeon** (Campagner et al., 2025) | Mouse | ✖ | ✔ | ✔ | ✖ | ✖ | ✔ | ✔ | ✔ |

**Table 1: Comparison between Training Village and other home cage automated systems available for rodents.** Columns indicate key features of each system: species tested, presence of an automatic training algorithm (✔ = yes, ✖ = no; 🔶 = not clearly specified), group housing capability, home-cage activity monitoring, isolated task testing determines whether animals are tested individually without interference from conspecifics, testing outside the home cage indicates whether the task is integrated within the home-cage or or externally connected (enabling greater modularity and flexibility), touchscreen availability, alarm system, and open-source software availability.

| Type | Name | Description | Urgency |
|---|---|---|---|
| Command | /report ‘hours’ | Provides a summary of the operant box activity for the last specified hours. | - |
| Command | /cam | Sends a screenshot from the two cameras. | - |
| Command | /plot ‘days’ | Shows a plot of entrances and attempts over the specified days | - |
| Report | report 24h | Automatically provides a summary of the operant box activity for the last 24h. | - |
| Alarm | Subject low water intake | The subject has consumed less than the configured water in the last 24 h. | Low* |
| Alarm | Subject no trials | The subject did not complete any trials in the behavioral session | Low |
| Alarm | Subject too long in box | More than an hour has passed since the task ended, and the animal remains inside the operant box. | Low |
| Alarm | No data sync | No data has been successfully synchronized with the external server or hard drive during the last 24 h. | Low |
| Alarm | Subject no sessions | The subject has not performed a behavioral session for the last 24 h. | Medium |
| Alarm | 2 Subjects in box | A high pixel count suggests multiple animals entered the operant box. | Medium |
| Alarm | Subject in a prohibited area | Pixels are detected in a region where no animal should be present. | High |
| Alarm | Subject no detected | The subject has not been detected by the RFID reader for the last 24 h. | High |
| Alarm | Error running task | Error running task. | High |
| Alarm | Temperature problems | The room temperature has dropped or exceeded the configured threshold in settings. | High |
| Alarm | Scale not responding | The scale is continuously returning a weight of 0.0 grams, indicating that it is not receiving valid readings from the load cell. | High |
| Alarm | No detection in 6h | No animal has been detected by the RFID reader in the last 6 h. | High |
| Alarm | Heartbeat not received | The system sends an hourly heartbeat to an external server; a missing heartbeat indicates a possible internet outage, power failure, or freeze. | High |

**Table 2: Summary of Telegram functions for remote supervision.** Commands can be used at any time to retrieve information from the system. Subject reports are displayed automatically twice per day (every 12 hours) and provide information about the general status of each animal in the system. Alarms are automatically triggered by the TV whenever a problem occurs. *Subject low water intake has low urgency if animals are provided with CA water in the home cage, otherwise, the urgency becomes high. Some critical alarms disable the RFID reader and halt new entries to protect other subjects.

| Group number | Specie | # Animals | Sex | Genotype | Task | Data collection (months) | Removed animals | Eco-HAB | # Home cages |
|---|---|---|---|---|---|---|---|---|---|
| 1 | Mouse | 8 | Male | 4 C57Bl/6J<br>4 Gad2-Cre | 3AFC | 1.5 | 0 | No | 2 |
| 2 | Mouse | 10 | Male | C57Bl/6J | 3AFC | 1 | 0 | No | 1 XL |
| 3 | Mouse | 10 | Male | C57Bl/6J | 3AFC | 1.5 | 0 | No | 1 XL |
| 4 | Mouse | 12 | Male | C57Bl/6J | 3AFC | 7 | 1 | Yes | 4 |
| 5 | Mouse | 12 | Female | C57Bl/6J | 3AFC | 6 | 1 | Yes | 4 |
| 6 | Mouse | 12 | Female | C57Bl/6J | 3AFC | 6 | 2 | Yes | 4 |
| 7 | Mouse | 14 | Male | 7 Pv-Cre<br>7 Pv-Cre Grin1 | 3AFC | 12 | 2 | Yes | 4 |
| 8 | Mouse | 14 | Male | C57Bl/6J | 3AFC | 2 | 0 | No | 2 XL |
| 9 | Rat | 6 | Female | Long Evans | 3AFC | 12 | 0 | No | 3 |
| 10 | Mouse | 3 | Male | C57Bl/6J | 2AB | 3 | 0 | No | 2 |
| 11 | Mouse | 10 | Male | C57Bl/6J | 2AB | 1.5 | 1 | No | 2 |
| 12 | Mouse | 10 | Male | TRAP2 x Ai14 | 2AFC | 1.5 | 0 | No | 2 |

**Table 3: Specifications of the different experimental groups.**
